# An Autonomous Microbial Sensor Enables Long-term Detection of TNT Explosive in Natural Soil

**DOI:** 10.1101/2024.10.08.617249

**Authors:** Erin A. Essington, Grace E. Vezeau, Daniel P. Cetnar, Emily Grandinette, Terrence H. Bell, Howard M. Salis

**Affiliations:** Department of Chemical Engineering, The Pennsylvania State University, University Park, PA, 16802; Department of Biological Engineering, The Pennsylvania State University, University Park, PA, 16802; Department of Biomedical Engineering, The Pennsylvania State University, University Park, PA, 16802; Department of Plant Pathology and Environmental Microbiology. The Pennsylvania State University, University Park, PA, 16802; Department of Physical and Environmental Sciences, University of Toronto.

## Abstract

Microbes can be engineered to detect target chemicals, but when they operate in real-world environments, it remains unclear how competition with natural microbes affect their performance over long time periods. We engineered sensors and memory-storing genetic circuits inside *Bacillus subtilis* to sense and respond to the TNT explosive, using predictive models for rational design. We characterized their ability to detect TNT in a natural soil system, measuring single-cell and population-level behavior over a 28-day period. The autonomous microbial sensor activated its response by 14-fold when exposed to low TNT concentrations and maintained stable activation for over 21 days, exhibiting exponential decay dynamics at the population-level with a half-life of about 5 days. Our results show that engineered soil bacteria can carry out long-term detection of an important chemical in natural soil with competitive growth dynamics serving as additional biocontainment.

## INTRODUCTION

Engineered microbes can sense target chemicals and respond with programmed actions, using cell sensors to detect specific chemicals and genetic circuits to convert regulatory signals into observable outcomes^1-3^. Microbial sensors have the potential to carry out autonomous, long-term environmental monitoring inside complex environments, such as soil systems or the human gut, without the need for direct observations or human intervention. Through such autonomy, microbial sensors can generate a rich stream of temporal and geospatial information that can be harnessed to dynamically guide human efforts, for example, fine-tuning fertilizer usage to avoid waste runoff, identifying geographic sources of environmental contamination, or detecting the presence of harmful pathogens^4-9^.

While this vision is enticing, autonomous microbial sensors need to maintain their functional performance for long periods of time (days to months) while operating in competitive environments, for example, surrounded by natural microbes that compete for scarce resources. It remains unclear how the design of the synthetic genetic circuit impacts the microbe’s ability to grow, persist, and perform in complex environments as resource constraints often result in synthetic genetic circuits that greatly inhibit cell growth or are lost via genetic instability^9-12^. To assess these effects, several types of quantitative measurements are needed to independently quantify persistence and function of autonomous microbial sensors over long time periods in operating environments. Engineering autonomous microbial sensors will require new design methodologies and multi-modal longitudinal test workflows.

One important autonomous microbial sensor application is the detection of the explosive compound 2,4,6-trinitrotoluene (TNT) inside natural soil systems that contain both engineered and naturally occurring microbes^13^. TNT was the predominant explosive used in ordinance from 1940-1970 and remains widely used in mines, grenades, antitank rockets, artillery, and bombs. Soil systems become contaminated with TNT from unexploded ordinance, leakage from deployed munitions, or from efflux during its manufacturing process. As TNT is insoluble in water, its release into soil systems can accumulate to high concentrations and results in a heterogeneous distribution of material with spatial hotspots^14^. While analytic techniques have been developed to detect TNT in soil samples, it is laborious and dangerous to carry out on-site sampling with sufficient geospatial resolution. Autonomous microbial sensors have the potential to provide geospatial data streams across large geographic areas.

In prior efforts, cell sensors have been developed to detect explosive compounds, utilizing either endogenous promoters, allosteric transcription factors, or engineered riboswitches to regulate gene expression. Genome-wide screens and promoter fusion experiments in *E. coli* have identified several natural promoters that respond to TNT^15, 16^, although likely using an indirect mechanism that depends on an unknown degradative product or non-specific activation of DNA damage repair pathways caused by TNT toxicity^17^. The XylR transcription factor from *Pseudomonas putida* was engineered via mutagenesis and DNA shuffling to improve its binding specificity to TNT, resulting in about 3-fold transcriptional activation at 1 mM TNT (227 mg/L) in liquid cultures^18^. Riboswitches have been engineered in *E. coli* to activate reporter protein expression by 4.6-fold in response to 0.1 mM TNT or by 11.1-fold in response to 1 mM DNT in liquid cultures^19, 20^, harnessing RNA aptamers for binding specificity^21^.

Here, we engineered the soil bacteria *Bacillus subtilis* to detect the explosive compound TNT inside a competitive soil system with a natural microbiome to carry out autonomous microbial sensing for over 3 weeks (**Figure 1A**). Our designed genetic circuit combines a TNT riboswitch sensor with a genetic memory switch, controlled by sense and antisense promoters, to stably activate an observable response function across long time periods (**Figure 1B**). We designed, built, and tested several autonomous microbial sensor variants, utilizing sequence-to-function biophysical models and system-wide models to rationally engineer synthetic genetic circuits for low-burden operation in *Bacillus subtilis*. We then quantitatively characterized the autonomous microbial sensor’s TNT sense-and-response function in a natural soil system across a 4-week longitudinal period, while measuring cell viability and persistence of the competing natural and engineered microbes. The best autonomous microbial sensor activated its response function by 14-fold after 1-week of TNT exposure (4.5 mg TNT/kg soil) with an exponential decay in viability and function over time, reaching 2.7-fold activation after 3 weeks of TNT exposure. We show that autonomous microbial sensors can survive long enough in complex environments to carry out a meaningful sense-and-response function, though our measurements suggest that natural microbes can serve as an effective source of biocontainment in competitive environments.

**Figure 1:**
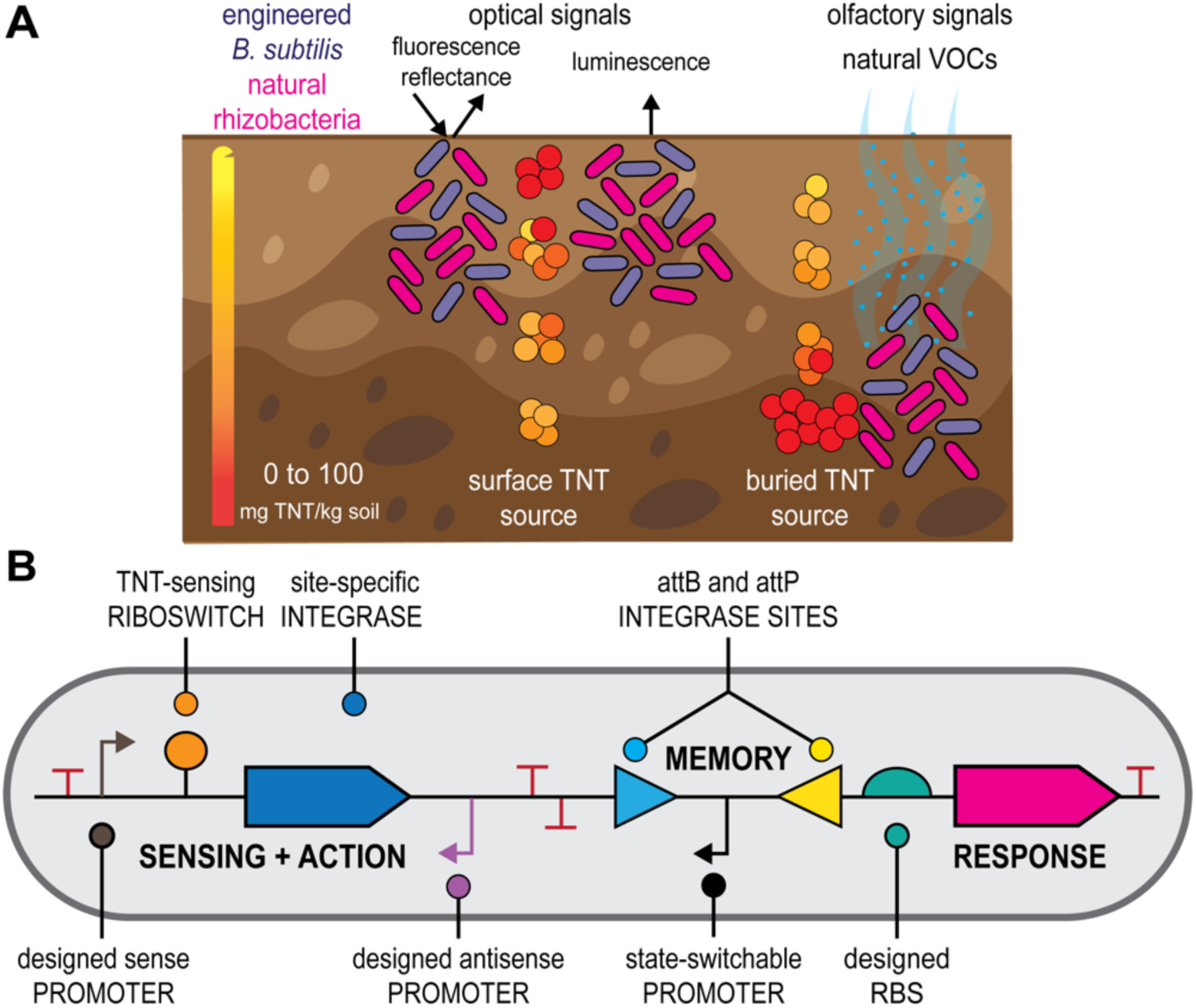
A synthetic genetic circuit for detecting TNT in soil. **(A)** Engineered *Bacillus subtilis* cells sensing TNT inside soil in competition with natural microbes, using optical or olfactory output signals to transmit response information for stand-off detection. **(B)** The engineered synthetic genetic circuit contains cell sensing, memory, and response modules. A constitutive promoter transcribes a TNT-sensing riboswitch, which activates the translation of a site-specific integrase. An antisense promoter reduces sensor leakiness via transcriptional interference. When expressed, the integrase binds to the attB/attP sites and flips the orientation of a promoter region. When flipped, the promoter expresses the output response module. Modules are insulated using transcriptional terminators.

## RESULTS

### TNT-sensing biosensor design and characterization at low TNT concentrations

We selected *Bacillus subtilis* 168 PS832 (Bacillus Genetic Stock Center) as a prototrophic *B. subtilis* strain able to grow autonomously in soil without amino acid supplementation, hereafter referred to as *B. subtilis*. We first attempted to identify and utilize natural promoters in *B. subtilis* that were transcriptionally regulated in response to TNT. We carried out transcriptome-wide mRNA level measurements (RNA-Seq) on *B. subtilis* cells grown in LB media with 0, 10 μM, or 66 μM TNT, followed by differential gene regulation analysis (**Methods**). From the RNA-Seq read counts, we identified several TNT-responsive genes with up to a 10-fold increase in mRNA levels when growing cells were exposed to 66 μM TNT (**Table S1**). We extracted promoter regions (130 to 215 bp long) controlling these genes’ expression levels and tested whether they activated a mRFP1 reporter’s expression level in response to TNT when utilized within a synthetic genetic circuit that was integrated into the *B. subtilis* genome. However, we found that these extracted promoter regions only activated mRFP1 expression by 1.4-fold with a dose-dependent response when exposed to up to 25 μM TNT during exponential growth (**Figure S1**).

The relatively low transcriptional activation from these promoters suggested that these sequences did not fully encode the TNT-responsive mechanism. For example, the extended promoter regions (500 bp) upstream of these genes are predicted to have multiple transcriptional start sites (**Figure S2**), each potentially regulated by different transcription factors. Based on the predominant start sites, the natural 5’ untranslated regions are also much longer (160-180 nt) than canonical ribosome binding sites, suggesting that regulatory RNAs may additionally regulate mRNA levels^22^. The TNT-responsive genes themselves are enriched in transporters (**Table S1**), indicating that TNT exposure could trigger a more indirect mechanism that may not be specific enough to utilize as a TNT biosensor. For these reasons, we decided to switch from using natural transcriptional regulation as a biosensor to engineering synthetic RNA-based TNT biosensors.

Next, we designed, constructed, and characterized TNT-sensing riboswitches that activate the expression of an output protein, harnessing a TNT-binding RNA aptamer as a direct sensing element^21^. We engineered two riboswitch sequences using the Riboswitch Calculator^19^, which is a statistical thermodynamic model that predicts and designs the sequence-structure-function relationship for translation-regulating riboswitches. The model calculates the ribosome-mRNA-ligand interactions that control the mRNA’s translation initiation rate and predicts how the translation rate changes when the aptamer binds to its target ligand at a particular ligand concentration, using the RBS Calculator v2.1 free energy model to calculate the ribosome’s binding free energy to mRNAs^23^. The Riboswitch Calculator was previously applied to successfully engineer riboswitches that bind to small molecules (theophylline, dopamine, thyroxine, fluoride, and 2,4-dinitrotoluene) as well as proteins (monomeric C-reactive protein, IL32γ)^24^. Here, we used the Riboswitch Calculator to design two riboswitches (RS14, RS15) that use a TNT-binding aptamer to activate mRFP1 expression in *B. subtilis* (**Methods**). In **Figure 2A**, we show the model-predicted mRNA structure of the RS14 riboswitch, illustrating the structural re-arrangements that are expected to take place when TNT binds to the aptamer domain. The Riboswitch Calculator predicts that the RS14 mRNA’s translation initiation rate will be low in the absence of TNT (242 au on the RBS Calculator v2.1 scale) and increase by 5.15-fold when 35 μM TNT is added. For the RS15 riboswitch, the model predicts a higher translation initiation rate in the absence of TNT (1121 au) with a 2.3-fold increase when 35 μM TNT is added (**Figure S3**). The multi-state ribosome-mRNA-ligand calculations and translation rate predictions are shown in **Figure S4** and **Figure S5**. All designed sequences and calculations are included in the **Supplementary Data**.

**Figure 2:**
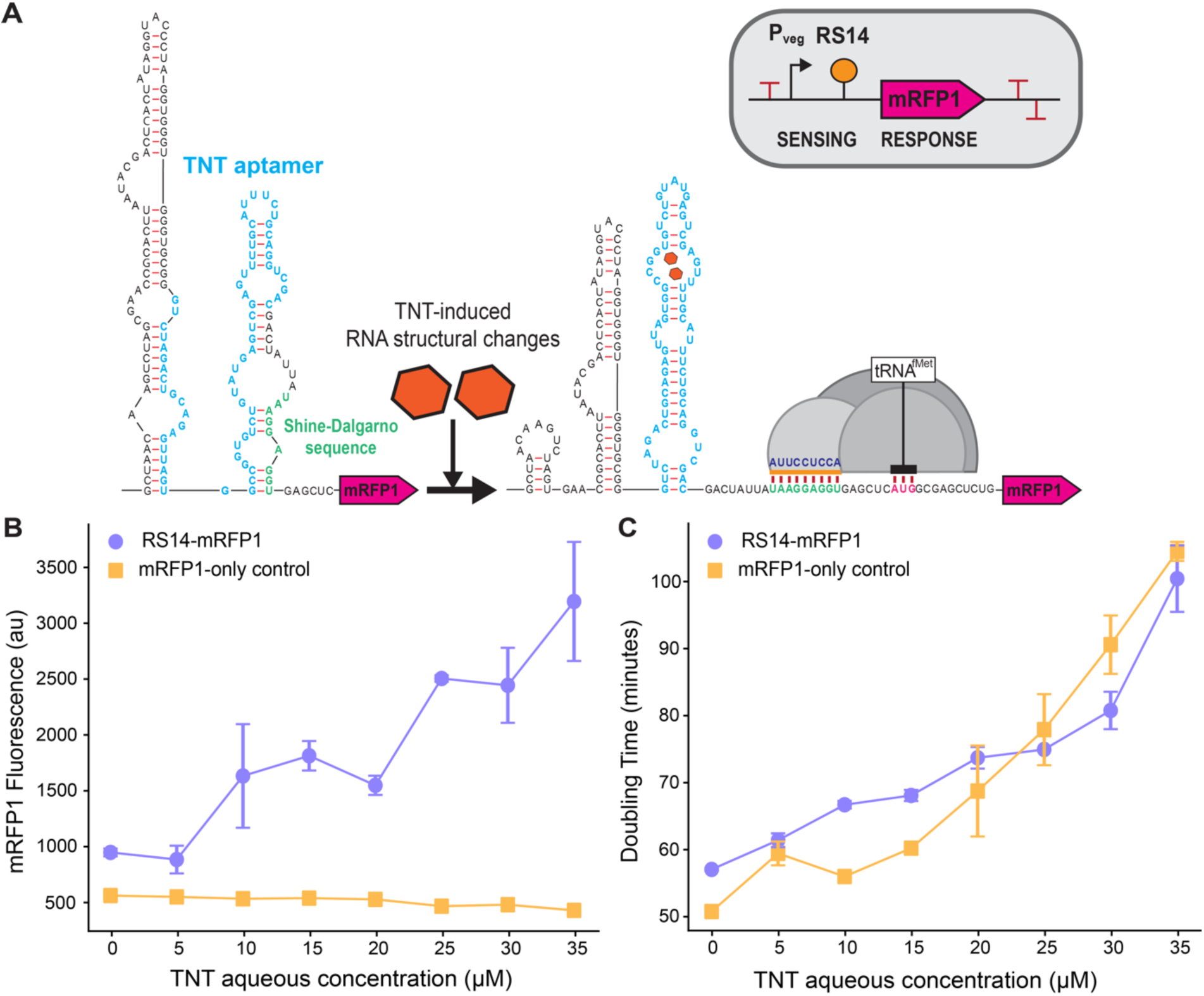
Design and characterization of a TNT-sensing riboswitch. **(A)** The Riboswitch Calculator model predicted mRNA structures of the RS14 riboswitch before and after induction with 35 µM of TNT. The “OFF” state (state 1) shows the mRNA structure and translation initiation rate calculated when TNT is unbound (rTL,OFF). The “ON” state (state 4) shows the change in mRNA structure and translation rate when both TNT is fully bound to its aptamer and the ribosome is bound to the mRNA (rTL,35mM). Nucleotides are color coded based on their interactions, (light blue) the TNT aptamer sequence, (green) the Shine-Dalgarno sequence (SD), (dark blue) the last 9 nucleotides of the 16S ribosomal RNA, and (pink) the start codon for mRFP1. The orange bar is the ribosomal footprint for initiation. For riboswitch characterizations, we added a strong *B. subtilis* promoter upstream to measure **(B)** the mRFP1 fluorescence levels and **(C)** the growth rates (doubling time in minutes) of the TNT-RS14 riboswitch strain (purple dots) and mRFP1-only control (yellow squares) in response to varied concentrations of TNT, up to 35 µM. Data points and error bars are the mean and standard deviation of N = 3 biological replicates.

We then constructed synthetic genetic circuits that use a P_veg_ promoter for constitutive transcription and either the RS14 or RS15 riboswitch to control the translation rate of a mRFP1 coding sequence. As controls, we constructed synthetic genetic circuits that constitutively express mRFP1 using a P_veg_ promoter and a designed ribosome binding site. All synthetic genetic circuits were integrated into the *amyE* locus of the *B. subtilis* genome using CmR for selection. We then cultured the engineered *B. subtilis* cells in M9 defined media supplemented with 2% glucose and with between 0 to 35 μM TNT added into aqueous solution, maintaining them in the exponential growth phase for at least 24 hours via serial dilution (**Methods**). By measuring their mRFP1 fluorescence levels, we found that both the RS14 and RS15 riboswitches activated mRFP1 expression in response to increased TNT concentrations (**Figure 2B, Figure S3**). The RS14 riboswitch increased mRFP1 fluorescence levels by 3.4-fold at 35 μM TNT with statistically significant activation (p-value = 0.018), while the RS15 riboswitch increased mRFP1 fluorescence levels by 2.1-fold in the same conditions (p-value = 0.006). In contrast, mRFP1 fluorescence levels from the mRFP1-only genetic circuit stayed largely constant and marginally decreased at the highest TNT concentrations (1.3-fold lower fluorescence, p-value = 0.009), likely due to TNT toxicity. Indeed, we measured cell doubling times and found that higher TNT concentrations caused *B. subtilis* cells to grow more slowly (**Figure 2C**, **Figure S3**). However, TNT toxicity impacted all synthetic genetic circuits similarly, whether they used a riboswitch to sense TNT or constitutively expressed mRFP1. Based on these results, we selected the higher performant RS14 riboswitch as a capable TNT biosensor.

### Engineering an autonomous microbial sensor for TNT detection

Soil systems are open environments with constantly changing concentration profiles. To achieve long-term detection of TNT in soil systems, our next step was to connect the TNT sensor to a genetic memory switch to convert transient TNT detection into stable activation of a desired response function. To do this, we designed and constructed new synthetic genetic circuits that use a constitutive sense promoter and the RS14 TNT-sensing riboswitch to regulate the expression of the Int2 serine integrase, followed by a constitutive antisense promoter that uses transcriptional interference to reduce basal (leaky) Int2 expression in the absence of TNT (**Figure 3A, Methods**). Int2 binds to a pair of recognition sites (attB and attP) and carries out site-specific recombination, flipping the orientation of DNA in between its recognition sites^25^. The next two components are a bidirectional transcriptional terminator to insulate circuit dynamics and a designed state-switchable promoter that is flanked by attB and attP sites. When the synthetic genetic circuit is built, the state-switchable promoter directs transcription in the reverse direction, but once Int2 expression is activated by the TNT sensor, its orientation will flip to the forward direction, triggering expression of the output response module. For quantitative circuit characterization, we selected the mRFP1 fluorescent reporter as an output response, using a designed ribosome binding site with a high translation rate (49000 au) so that the circuit behavior is readily observable even when integrated into the *Bacillus subtilis* genome. Overall, when the riboswitch binds to TNT, it activates Int2 expression, which flips the state-switchable promoter’s orientation and activates mRFP1 expression.

**Figure 3:**
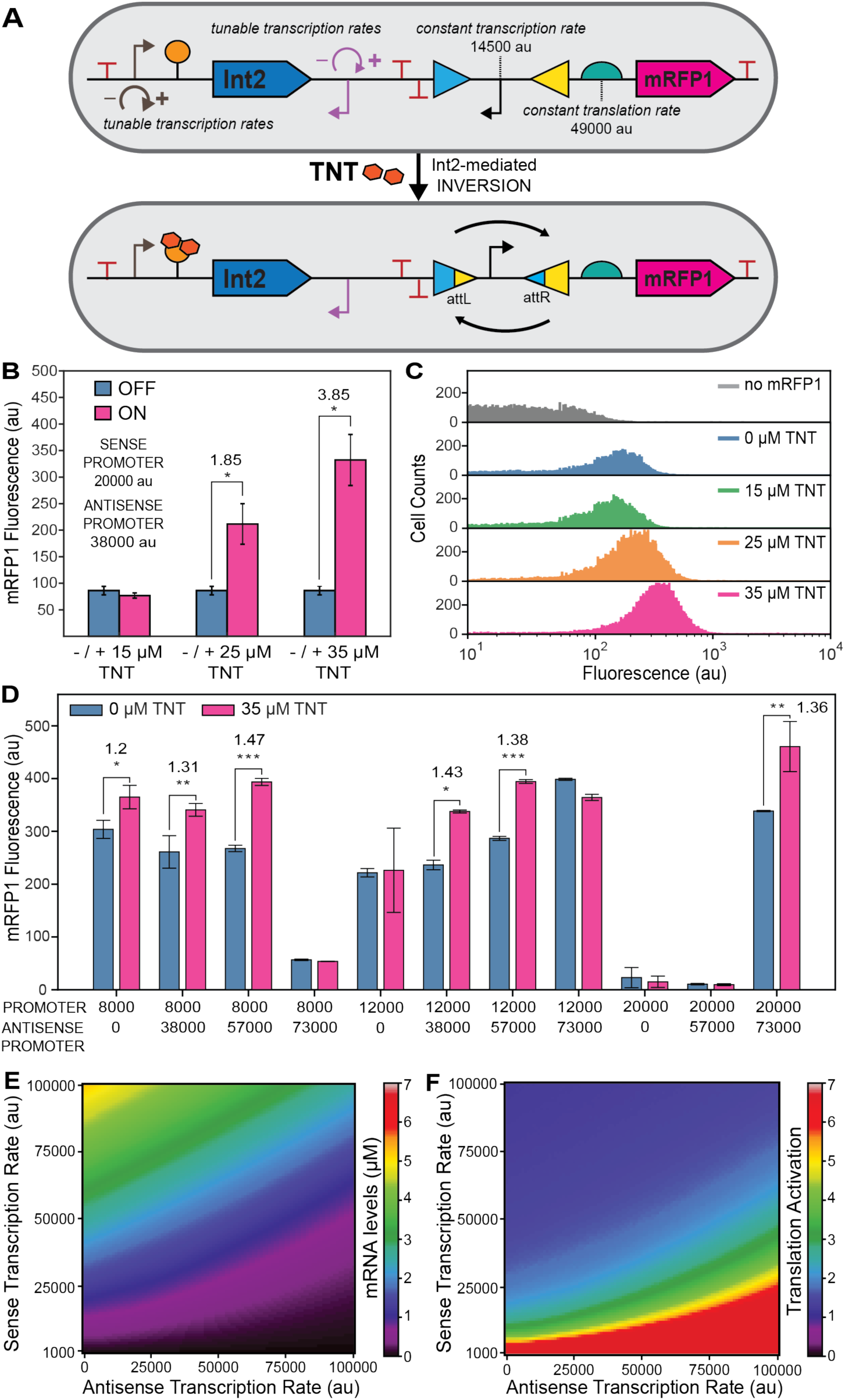
Design and testing of autonomous microbial TNT sensors in liquid culture. **(A)** TNT activates the synthetic genetic circuit by binding to the RS14 riboswitch, which activates expression of the Int2 recombinase. The Int2 recombinase binds to the att recognition sites and flips the state-switchable promoter’s orientation, which activates expression of a mRFP1 reporter (output module). Designed sense promoters control the transcription rate of the mRNA encoding the RS14 riboswitch and Int2 recombinase. Designed anti-sense promoters control the reduction in mRNA levels, due to transcriptional interference. Numbers are predicted transcription and translation rates. **(B)** Average mRFP1 fluorescence levels of engineered *B. subtilis* cells during exponential growth in liquid culture carrying the most performant synthetic genetic circuit in response to varied TNT concentrations (0, 15, 25, and 35 µM). Bars and error bars are the mean and standard deviation of 3 biological replicates. Numbers are activation ratios with statistical significance (p-values less than *0.05, **0.01, ***0.001). **(C)** The single-cell mRFP1 fluorescence distributions are shown for these engineered *B. subtilis* cells responding to each TNT concentration versus a wild-type control (no mRFP1). **(D)** Average mRFP1 fluorescence levels of engineered *B. subtilis cells* carrying different synthetic genetic circuits growing in the same conditions, comparing responses at 0 µM TNT (blue bars) versus responses at 35 µM TNT (red bars). Bars and error bars are the mean and standard deviation of 3 biological replicates. Quantitative models of **(E)** transcriptional interference and **(F)** riboswitch regulation show how changes in sense promoter and antisense promoter transcription rates alter Int2 mRNA levels and translation activation.

Through genetic fine tuning, we found that the transcription rates of the constitutive sense and antisense promoters controlled the synthetic genetic circuit’s input-output performance by modulating the dynamic range of Int2 expression across different TNT concentrations (**Figure 3BCD**). We rationally controlled these transcription rates by applying the Promoter Calculator^26^ – a thermodynamic model that predicts the site-specific transcription initiation rates of bacterial DNA sequences – to design sense and antisense promoter sequences with targeted rates from 8000 to 73000 on a proportional scale. For comparison, the P_veg_ promoter commonly used in *B. subtilis* has a predicted transcription initiation rate of about 38000 on the same scale. With this approach, we designed and constructed several synthetic genetic circuit variants, introducing combinations of sense and anti-sense promoters with varied transcription rates, followed by integrating them into the *B. subtilis* genome and measuring their output responses when cells were maintained in the exponential growth phase in defined M9 media supplemented with 2% glucose and between 0 to 35 μM TNT in aqueous solution (**Methods**). As part of our design criteria, we purposefully do not maximally over-produce the output response, both to reduce leaky expression in the absence of TNT as well as to reduce the overall cellular burden when responding to TNT.

The most performant synthetic genetic circuit activated mRFP1 fluorescence levels by 3.85-fold at 35 μM TNT, using a sense promoter (20000 au) and antisense promoter (38000 au) to optimally control transcription of the RS14 riboswitch and Int2 recombinase with circuit activation beginning to take place at around 25 μM TNT (**Figure 3B**). We found that increasing the TNT concentration caused a gradual activation of the synthetic genetic circuit’s response with a continual rightward shift of the single-cell mRFP1 fluorescence distribution (**Figure 3C**). We also applied reverse transcription qPCR (RT-qPCR) to directly measure mRFP1 mRNA levels after only 5 minutes of exposure to 35 μM TNT and found that they increased by 2.95-fold as compared to a no-TNT control (**Figure S6**). We then tested whether the synthetic genetic circuit could stably maintain a high output response after TNT was no longer present in the system. After growing the engineered *B. subtilis cells* in liquid culture with 35 μM TNT, we serially diluted into fresh media without TNT and found that mRFP1 fluorescence remained high (3.78-fold activation) for over 8 hours of additional exponential growth (**Figure S7**). These additional experiments show that the synthetic genetic circuit rapidly responded to TNT exposure, triggering Int2 recombinase expression activation, flipping the promoter DNA, and activating transcription of the output response module. After TNT is removed from the system, the state-switchable promoter remained in the flipped state and maintained stable expression of the output response module in actively growing cells.

We then carried out response characterization on all synthetic genetic circuits and found that excessively high or low transcription rates resulted in lowered genetic circuit activation ratios (**Figure 3D**). For example, when only a sense promoter was used to express the TNT riboswitch sensor controlling Int2 expression, genetic circuit activation was poor (1.2-fold) at the lowest transcription rate and non-existent as sense transcription rates were increased. However, very high antisense promoter transcription rates ablated genetic circuit activation, particularly when sense transcription rates were low. We were intrigued by this non-linear relationship between specification and circuit performance, motivating the development of system-wide models that accounts for known gene regulatory mechanisms to ascertain whether they could explain our observations.

We began by positing that three mechanisms related to transcriptional interference dynamics, molecular crowding, and recombinase toxicity were responsible for this non-linear behavior. First, when an antisense promoter is positioned downstream of a sense promoter, it lowers the sense mRNA levels via RNA polymerase collisions as well as degradation of the produced double-stranded RNA^27^. We derived and solved a system of partial differential equations that quantifies these mechanisms to calculate how changing transcription initiation rates at the sense and anti-sense promoters alters mRNA levels (**Figure 3E**, **Supplementary Information**). Second, increasing the TNT riboswitch sensor’s mRNA level is expected to increase Int2 expression, up to a point; however, as mRNA levels increase, the fold-activation in response to TNT can actually decrease as greater amounts of mRNA compete for binding to a limited amount of TNT ligand inside a crowded cellular environment. This phenomenon was previously observed across riboswitches that bind to diverse ligands^19^. To quantify this interaction, we applied the Riboswitch Calculator model to calculate how changing the TNT riboswitch’s mRNA level alters the fraction of riboswitch mRNA bound by TNT ligand and its corresponding Int2 translation rate. By using the solution to the first model as the input into the second model, our calculations show that Int2 translation activation is highest when the antisense promoter transcription rate is higher than the sense promoter transcription rate (**Figure 3F**, **Supplementary Information**). Finally, over-expression of the Int2 recombinase was found to be cytotoxic, creating an upper viability limit to the Int2 mRNA level. Altogether, the models show the trade-off between sensing TNT and achieving high Int2 expression activation, while ensuring that activation produces enough Int2 to flip the state-switchable promoter. Based on these models, the optimal design specification is a sense promoter with a moderately high transcription rate together with an antisense promoter that has a higher-than-sense transcription rate, which is consistent with our measurements.

### Long-term testing of an autonomous microbial sensor in natural soil

We developed an experimental workflow to quantitatively measure the persistence, viability, and functional response of a TNT-sensing autonomous microbial sensor in a soil system with a natural microbiome over a 28-day period. For these measurements, we selected the engineered *B. subtilis* strain carrying the most performant synthetic genetic circuit (**Figure 3B**). We first collected natural soil from an agricultural field and transferred 35 grams soil into containers at a depth of about 1 cm, followed by supplementation with a nutrient mixture containing ribose and hexose sugar (6 g/kg soil) and urea (0.1 g/kg soil) (**Methods**). In prior experiments, we found that the nutrient mixture increased the persistence of engineered *B. subtilis* cells by about 10-fold (**Figure S8**). We then evenly dispersed 2 mL of a TNT/DMSO solution to each container, using solutions with different TNT concentrations to vary the in-soil TNT amount from 0 to 110 mg TNT per kg soil. The DMSO was evaporated by incubation at room temperature, followed by even dispersal of about 10^8^ colony forming units (CFUs) of engineered *B. subtilis* per gram soil. Containers were then incubated at 25°C with periodic watering every 3 to 4 days to maintain hydration at 80-85% of saturation, using wrapping with parafilm to limit desiccation. We then extracted 500 mg soil samples every 3 to 4 days. We carried out several types of measurements on soil samples, including flow cytometry to measure single-cell mRFP1 fluorescence level distributions, sample plating to measure the number of living cells as quantified by the number of CFUs, quantitative PCR (qPCR) to measure the number of engineered *B. subtilis* genome regions, and next-generation sequencing to determine the genetic stability of the synthetic genetic circuit (**Figure 4A**).

**Figure 4.**
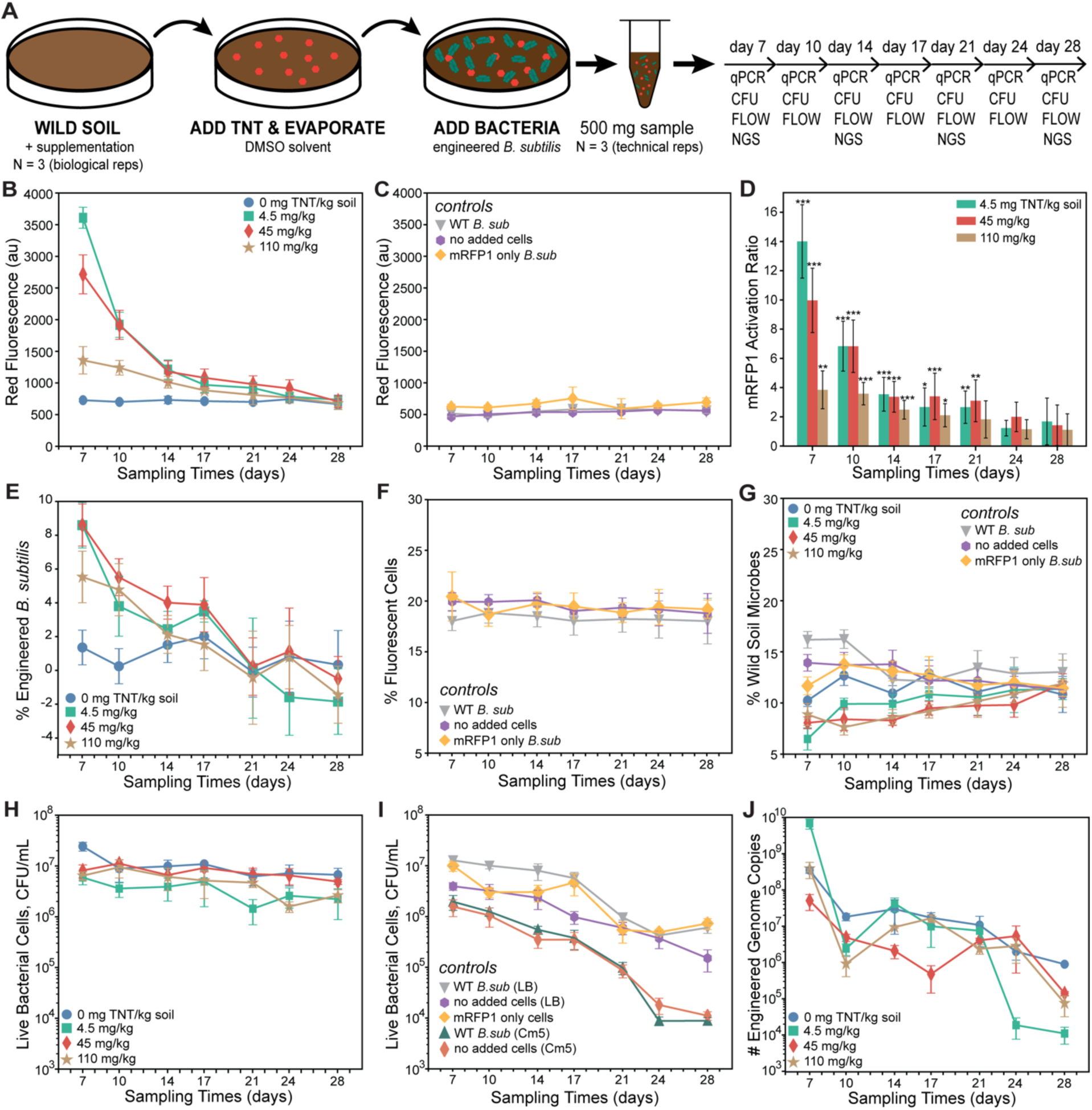
Long-term testing of an autonomous microbial TNT sensor in wild soil. **(A)** The experimental workflow and measurement timelines for characterizing the function and persistence of the TNT-sensing autonomous microbial sensor in wild soil. Measurements on soil samples included quantitative PCR (qPCR), colony forming units (CFUs), flow cytometry (FLOW), and next-generation sequencing (NGS). **(B)** Measured red fluorescence levels of the TNT-sensing autonomous microbial in wild soil at 0 mg TNT / kg soil (blue circles), 4.5 mg TNT / kg soil (green squares), 45 mg TNT / kg soil (red diamonds), and 110 mg TNT / kg soil (tan stars) over a 28-day period. **(C)** Measured red fluorescence levels of WT *B. subtilis* cells (grey triangles), mRFP1 only *B. subtilis* cells (orange diamonds), and the no added cells control (purple octagons) in wild soil without TNT over a 28-day period. **(D)** Measured mRFP1 activation ratios of the TNT-sensing autonomous microbial sensor in wild soil with 4.5 mg TNT / kg soil (green bars), 45 mg TNT / kg soil (red bars), and 110 mg TNT / kg soil (tan bars) over a 28-day period. The statistical significance of the activation ratios are shown (two-tailed T-test p-values less than * 0.05, ** 0.01, *** 0.001, N = 3). **(E)** The percentage of autonomous microbial sensor in wild soil with varied TNT amounts over a 28-day period. **(F)** The percentage of auto-fluorescent control cells in wild soil without TNT over a 28-day period. **(G)** The percentage of other natural microbial cells in wild soil with varied conditions across a 28-day period. **(H)** Measured cell viability counts on selective agar plates (LB/Cm5) from wild soil samples containing the autonomous microbial sensor with varied TNT amounts. **(I)** Measured cell viability counts on selective (LB/Cm5) or non-selective (LB) agar plates from wild soil samples containing controls. **(J)** Measured copy number of autonomous microbial sensor genomes in wild soil with varied TNT amounts as quantified by qPCR. Data points and error bars are the mean and standard deviation of **(B-I)** 3 or **(J)** 2 biological replicates (independent soil containers), each with **(B-I)** 3 or **(J)** 2 technical replicates (samples per container).

In addition to the TNT-sensing autonomous microbial sensor, we carried out the same experimental workflow on engineered *B. subtilis* strains carrying different synthetic genetic circuits as well as other controls. For comparative analysis, we inoculated soil containers with engineered *B. subtilis* strains that only used the TNT riboswitch sensor to activate mRFP1 reporter expression (“sensor only”) to determine how the memory switch controlled the sensor output and its dynamics over time (**Figure S9**). We also tested an engineered *B. subtilis* strain that constitutively expressed the mRFP1 reporter (“mRFP1 only”) at a detectable level to determine how a low, constant expression burden affected the strain’s competitiveness (**Figure S10**). For another comparison, we inoculated the soil containers with wild-type (WT) *B. subtilis* cells as a baseline comparator. Finally, we carried out the same measurements on soil containers that were not inoculated with any additional cells, leaving only the natural microbes. All measurements, half-life calculations, statistical analyses can be found in **Supplementary Data**.

Overall, we found that the TNT-sensing autonomous microbial sensor achieved 14-fold activation of its output module after 7 days of low TNT exposure (4.5 mg TNT/kg soil) in natural soil, though there are important environmental and dynamical processes occurring simultaneously that affect the microbial sensor’s function and persistence as it competes with the soil microbiome (**Figure 4**). First, we observed an exponential reduction in single-cell fluorescence levels (sensor output) over the 28-day period with an apparent half-life of 5.4 to 7.9 days (**Supplementary Data** for analysis), depending on the amount of TNT added to the soil (**Figure 4B**). As more TNT was added to the soil (45 and 110 mg TNT/kg soil), cell function became inhibited and the sensor output activation was 10-fold and 4-fold, respectively, after 7 days of exposure (**Figure 4D**). After 21 days, the autonomous microbial sensor maintained an output activation of 2 to 3-fold (**Figure 4D**). Interestingly, our engineered “sensor only” *B. subtilis* strain also activated mRFP1 expression, but at a lower magnitude (2.4-fold at day 7), and with a similar exponential reduction over time (half-life of 12 days) (**Figure S8**). In contrast, our engineered “mRFP1 only” *B. subtilis* cells produced a low, but detectable, amount of fluorescence, similar to the autonomous microbial sensor’s output in the absence of TNT (**Figure 4C**). We also found that auto-fluorescence levels from the wild-type *B. subtilis* cells and from natural microbes already in the soil were relatively constant over time (**Figure 4C**).

We then quantified how the relative proportion of natural versus engineered microbes changed over time and across varying amounts of TNT added to the wild soil, using flow cytometry to identify two key sub-populations of cells (**Methods**). The first cell sub-population contained a mixture of autonomous microbial sensors and natural microbes, which generally had larger size characteristics, while the second cell sub-population were only natural microbes that had generally smaller size characteristics (**Figure S11**). We identified the engineered *B. subtilis* cells within the first cell sub-population based on their higher red fluorescence levels, as compared to the controls. From these measurements, we determined the proportion of autonomous microbial sensors in the first cell sub-population, which we then tracked over time (**Figure 4E**). We found that adding TNT into the wild soil caused the autonomous microbial sensors to begin at a relatively high proportion (6-8%), followed by an exponential reduction over time (half-lives varied from 4.1 to 6.5 days), reaching the control baseline after 24 to 28 days. In contrast, when TNT was not added to the soil, the autonomous microbial sensor and control cells both maintained a relatively constant proportion (**Figure 4F**). By tracking the dynamics of the second cell sub-population, containing only natural microbes, we found that adding TNT did initially lower their proportion (7-9%), but they largely rebounded and increased in proportion over time, reaching about 10-12% after 28 days (**Figure 4G**). As a comparison, the proportion of natural microbes in the control experiments stayed largely constant over the same time period (**Figure 4G**). The remaining percentage of unlabeled events stayed largely constant over time and across TNT conditions, suggesting that they are soil debris. Overall, the engineered cells were able to initially exist as a significant minority proportion of the total cell population, but over time, the natural cells proliferated and eventually overtook the engineered ones.

We also carried out cell viability assays on all samples to quantify the number of live cells according to their colony forming units on selective or non-selective LB agar plates (**Methods**). The number of living autonomous microbial sensor cells in wild soil stayed stable over time, decreasing by only 1.2 to 3.7-fold over the 28-day period (half-life of 14 days), depending on the amount of TNT exposure (**Figure 4H**). However, the addition of TNT to the soil had an overall cytotoxic effect; adding 110 mg/kg TNT lowered CFU counts by 1.3 to 4.5-fold across all timepoints. In comparison, the persistence of wild-type *B. subtilis* in a soil system without TNT exposure was less stable (half-life of 3.9 days), which was similar to the persistence of the wild microbes in the same condition (half-life of 4.2 days) (**Figure 4I**). As expected, about 25% of the natural microbes in these soil systems were naturally resistant to low antibiotic concentrations (Cm5).

Interestingly, from the qPCR measurements, we found larger reductions in the engineered genome’s copy number over the 28 day period (**Figure 4J**), though the qPCR measurements had far higher variability across samples and conditions as compared to the live cell counts. The engineered DNA can originate from live or dead cells, and remains subject to chemical changes over the 28 day period, which could explain why the qPCR measurements exhibited higher variabilities.

Finally, we investigated the genetic stability of the engineered genetic circuit over the 28-day period. To do this, we extracted genomic DNA from the soil samples taken at days 7, 14, 21, and 28, and then carried out PCR to selectively amplify the circuit region that was integrated inside the engineered *B. subtilis* genome, followed by long-read nanopore sequencing (**Methods**). Perhaps surprisingly, we did not find any DNA mutations in the genetic circuit over the 28 day period. Notably, the genetic circuit was integrated into the *B. subtilis* genome and was designed to have few repetitive DNA sequences with a maximum DNA repeat length of only 12 base pairs, which together eliminate several sources of DNA recombination and lower the rate of evolutionary selection.

## Discussion

We engineered a synthetic genetic circuit in *Bacillus subtilis* capable of long-term detection of TNT and characterized its ability to function and persist while competitively growing in wild soil with a natural soil microbiome. The autonomous microbial sensor achieved 14-fold output activation after 7 days of TNT exposure at a relevant contaminant concentration of 4.5 mg TNT/kg soil, while maintaining measurable activation of its output response module for 21 days, which altogether is a sufficiently long time to use area-wide measurements to survey large geographic areas. We found that the number of engineered *B. subtilis* cells and their output module activation both exponentially decayed over the 28 day testing period with half-lives of about 4 to 8 days, depending on the amount of TNT added to the contaminated soil. These exponential decay dynamics indicate that the engineered cell population diminishes to 1% of its initial number within about 27 to 53 days. Altogether, our results show that autonomous microbial sensors can function long enough in a microbially competitive natural environment to provide valuable geospatial information, answering a long-standing question in the field.

However, careful genetic system design and optimization was needed to obtain long-term performance. Overall, it was important to design the genetic circuit to avoid over-expressing components, both to lower the output response in the absence of TNT (leakiness) and to reduce the overall expression of the genetic circuit’s components (cellular burden). It was also important to ensure the absence of repetitive DNA sequences in the genetic circuit and to integrate the genetic circuit into the *Bacillus subtilis* genome to improve the evolutionary stability of the genetic circuit and the resilience of the organism’s response. These design criteria lower the probability of introducing circuit-breaking mutations and reduce the selectiveness of the fitness landscape, particularly in *B. subtilis* where the intrinsic rate of homologous recombination is relatively high.

To carry out and test our designs, we first applied statistical thermodynamics and computational optimization (the Riboswitch Calculator^19^) to rationally engineer TNT-binding riboswitch sensors with well-characterization dosage response curves in both liquid culture and natural soil environments (**Figure 2**). We then embedded the TNT sensor within a tunable synthetic genetic circuit whose purpose is to convert transient signal detection into stable activation of a modular output response, using translation-activation of a DNA-flipping integrase to irreversibly switch the circuit from an OFF to ON state (**Figure 3**). To fine-tune the circuit input-output response and lower leaky activation in the absence of TNT, we used thermodynamic modeling to design sense and anti-sense promoters with desired transcription initiation rates, using the Promoter Calculator^26^. We developed system-wide gene regulation models to guide our design-build-test cycling and to understand how changing the promoters’ transcription rates altered sensor and circuit function. We then carried out longitudinal testing of the best TNT-sensing autonomous microbial sensor in natural agricultural soil for a 28 day period, utilizing flow cytometry, qPCR, colony counting, and next-generation sequencing to quantitatively measure single-cell and population-level characteristics of both engineered and natural microbes (**Figure 4**). Such measurements are needed to determine the longevity of sensor-circuit function in a perturbed real-world environment as well as the long-term effects of the engineered bacteria on the natural microbial population.

The synthetic genetic circuit was designed to be modular and tunable with the potential to readily introduce new sensors to detect target stimuli. Cell sensors utilizing inducible transcription factors or two-component systems can be used to activate or de-repress transcription of the integrase in response to a wide variety of chemicals, including oxygen, nitrates, phosphates, excess metals, insecticides, quorum sensing signals, and toxins^2, 3,8^. Novel riboswitches can be engineered to activate translation of the Int2 integrase in response to additional chemicals that may bind poorly to proteins, such as pesticides^28, 29^, heavy metals^30^, and perfluorooctanoic acids (PFOAS)^31^, utilizing model-based design algorithms (e.g. the Riboswitch Calculator) to rationally convert aptamers into high-performance riboswitches. Oftentimes, cell sensors exhibit different amounts of leaky expression even in the absence of the target stimuli, however, our synthetic genetic circuit uses a tunable anti-sense promoter to reduce leaky expression via transcriptional interference, which increases the sensor’s dynamic range.

The synthetic genetic circuit’s architecture also decouples the sensor and output modules, enabling the introduction of new output modalities without having to modify the cell sensor. Here, we used the mRFP1 fluorescent protein reporter for more precise quantitative measurements in liquid culture and soil systems. For line-of-sight geospatial measurements, it would be advantageous to introduce a single or multi-enzyme pathway to produce chemically stable pigments with readily detectable hyperspectral emission wavelengths, such as carotenoids and melanin-containing biopolymers. However, when it becomes important to sense target chemicals without line-of-sight detection, for example, detecting TNT from a buried munition, new output modalities will be needed that transmit information via chemical signals that can quickly diffuse through its environment and bind to other cell sensors, including animal olfactory systems. Finally, rather simply detecting and responding to TNT, the output module could additionally include the expression of nitroreductase enzymes that degrade TNT into non-toxic compounds, creating an integrated sensing and bioremediation capability.

As TNT soil contamination remains a world-wide and ongoing challenge, autonomous microbial sensors are a promising solution to detecting TNT hotspots across large geographic areas without on-site sampling. While there are nation-specific regulatory requirements that determine when a genetically modified organism can be deployed in an outdoor environment, it remains unclear how the organism’s behavior in an ecosystem should affect these regulatory decisions. Our results show that the competition between engineered and natural microbes can be well-characterized at the population-level, quantifying the need for biocontainment at a desired time-scale. Thus, as part of an organism engineering pipeline, autonomous microbial sensors can be developed to sense a target chemical, respond with a desired program, and disappear from their environment with a reproducible timeline. A scenario-specific, data-driven approach would greatly refine and clarify regulatory requirements, enabling the deployment of well-tested autonomous microbial sensors that see the unseen world around us.

## Methods

### Design of TNT-sensing riboswitch sequences using Riboswitch Calculator

We obtained the TNT aptamer sequence and binding affinity (K_D_ = 1×10^-8^ M) from published literature^21^. The aptamer’s secondary structure was determined using RNAfold (Vienna RNAfold v2.5)^32^. We used the design mode of the Riboswitch Calculator to computationally optimize pre-aptamer and post-aptamer sequences that are predicted to maximize the mRNA’s translation initiation rate when the RNA aptamer is bound by TNT, while minimizing the mRNA’s translation initiation rate when the RNA aptamer is not bound by TNT^19, 24^. The design inputs include the RNA aptamer sequence, the structure of the RNA aptamer when bound by TNT, the ligand-aptamer binding affinity, and the mRFP1 coding sequence whose expression level is being regulated by the riboswitch. From the output list of designed riboswitch sequences, we selected two sequences (RS14 and RS15) based on their predicted maximum activation ratios and translation rates in the off and on states. The predicted mRNA structures of riboswitches were partly made using Forna diagrams^32^.

### Whole Transcriptome RNA-seq Quantification and Promoter Extraction

The *Bacillus subtilis substr*. 168 PS832 strain was initially cultured for 12 hours in LB media, followed by serial dilutions into M9 minimal media supplemented with 0, 10, or 66 μM TNT, maintaining cells in the exponential growth phase for at least 16 hours. Total RNA was extracted from samples using Total RNA Purification Kit (Norgen Biotek), followed by DNA removal using the Turbo DNAse Kit (Ambion). Purified RNA samples were then rRNA depleted using the NEBNext rRNA Depletion Kit (NEB), followed by cDNA synthesis using SuperScript IV First-Strand Synthesis System (Invitrogen) and random hexamer primers, followed by second strand synthesis (Q5 DNA polymerase). NGS library preparation and RNA-Seq were carried out by GeneWiz, yielding over 50 million reads per triplicate sample. For RNA-seq analysis, a *k*-mer filtering approach (BBDuk) was used to remove reads containing rRNA or noncoding RNA, using the *B. subtilis substr.* 168 PS832 RefSeq genome (NC_000964). Reads containing a 31-mer match (with up to one mismatch) were removed. The remaining RNA reads were mapped and aligned to CDS features. Transcriptome abundances were aligned using HISAT2 v2.1.0 and counted using featureCounts to obtain average TPM measurements for each CDS. The R package edgeR (v.3.22.5) was used to calculate the differential expression of genes from the HISAT2-aligned reads, using default settings. Genes were selected if they were significantly differentially expressed (*P*<0.05), resulting in identification of at least five TNT-responsive promoters that had up to a 9.97-fold change in RNA-seq read counts (TPMs) when exposed to 66 μM of TNT (Supplementary Data).

### Plasmid Design and Cloning

Integration vectors were prepared using the vector fragment of pDG1662, which was obtained from Bacillus Genetic Stock Center. The vector fragment contains a ColE1 origin and two amyE’ homology arms that surround the insertion site for designed systems and a CmR marker for selection of genome integration events. To prepare the vector fragment, an inverse PCR reaction was carried out on pDG1662 using primers pDG1662_rev_1 and pDG1662_fwd_2. Designed genetic systems were synthesized as gBlock DNA fragments (Integrated DNA Technologies), PCR amplified, and assembled with the vector fragment using HiFi assembly (New England Biolabs). All genetic part sequences and primers are included in the Supplementary Data. During the design of genetic systems, all transcription initiation rate predictions were carried out using the Promoter Calculator v1.0 model and all translation initiation rate predictions were carried out using the RBS Calculator v2.1 model. All protein coding sequences were codon optimized for expression in *Bacillus subtilis*, using the CDS Calculator to remove any repetitive DNA, internal promoters, highly translated internal start codons, or unique cut sites.

To test TNT riboswitch sensors, genetic systems were designed to contain a constitutive *B. subtilis* promoter (P_veg_, 30,000 au on the Promoter Calculator scale), either the RS14 or RS15 riboswitch, a mRFP1 reporter coding sequence, and flanking PCR primer binding sites. As a control, a mRFP1-only genetic system was designed to contain the P_veg_ promoter, a designed ribosome binding site (20,000 au on the RBS Calculator scale), a mRFP1 coding sequence, and the same flanking PCR primer binding sites. To test TNT inducible promoters, genetic systems were designed to contain natural *B. subtilis* promoter regions (QX56-11955 or QX56-01665), followed by a designed ribosome binding site (25000 au), a mRFP1 reporter coding sequence, and flanked by the same PCR primer binding sites.

To test genetic circuit variants, genetic systems were designed to contain a constitutive promoter, the RS14 TNT-sensing riboswitch, a site-specific integrase (Int2), tandem transcriptional terminators (ECK120017009 and ECK120010850), the integrase-specific recognition sites (attB and attP) flanking a state-switchable constitutive promoter (500 to 15000 au), a ribosome binding site (25,000 au), and a mRFP1 coding sequence.

For the data set Circuit Variant Testing, a baseline cassette was used and inserted into the pDG1662 cloning vector. The expression cassette included a constitutive natural promoter (P_veg_), a TNT-sensing riboswitch (RS14), a site-specific integrase (Int2), strong double terminators (ECK120017009 and ECK120010850), the integrase-specific recognition sites (attB and attP) harboring a state-switchable promoter (P501 – 15,000 au), a strong RBS (25,000 au), and the mRFP1 fluorescent protein. The cassette was flanked by primer binding sites, designed for PCR amplification and insertion into the vector by multi-fragment assembly. Unique restriction sites were placed throughout the cassette to facilitate the replacement of specific genetic parts.

The upstream constitutive promoters tested were assembled by designing individual gBlocks that contained the desired promoter upstream of the riboswitch, flanked by a 5’-BamHI and 3’-SbfI restriction site and primer binding sites. New promoters were inserted into the plasmid using digestion and ligation. The antisense promoters tested were assembled by annealing two complementary oligonucleotides with 5’FseI and 3’-SpeI restriction overhangs. The annealed oligonucleotides were ligated into the desired plasmid downstream of the integrase and upstream of the double terminators, either replacing or adding an antisense promoter.

All constructed plasmids were transformed into chemically competent *E. coli* DH10B cells and the isolated clones were sequenced verified (Quintara Biosciences and Plasmidsaurus). Miniprepped plasmids with correct DNA sequences were then genome integrated into *Bacillus subtilis* substr. 168 PS382 (Bacillus Genetic Stock Center Catalog: *Bacillus subtilis* 168, #1A757) for testing.

### Bacillus subtilis Integration

All plasmids were integrated into the genome of *Bacillus subtilis* substr. 168 PS832 using the following protocol. A single colony from a freshly streaked plate was selected and inoculated into 10 mL of MD medium (5 mL of 10X PC solution, 5 mL of 20% (v/v) glucose, 2.5 mL of 50 mg/mL DL aspartic acid potassium salt, 250 μL of 2.2 mg/mL ammonium iron (III) citrate, and 150 μL of 1M MgSO4) with the addition of 50 μL of 20% casamino acids. Cultures were incubated at 37°C and 300 RPM orbital shaking until an OD600 of 1-1.5 was reached. Once at the proper OD_600_, a 10 mL aliquot of MD medium, with no casamino acids, was added to the culture and incubated again at 37°C and 300 RPM orbital shaking for 1 hour. Purified plasmid DNA was then added to 800 μL of the now competent *B. subtilis* cells at a final concentration of 1 μg/mL and incubated for 20 minutes at 37°C and 300RPM orbital shaking. Finally, 25 μL of 20% casamino acids was added to the culture and incubated for 1 hour at 37°C and 300RPM orbital shaking. Cells were plated on chloramphenicol (5 μg/mL) and 1% starch agar plates and incubated at 30°C for 16-20 hours. To screen the colonies for disruption of the amyE’ locus (successful integration), a Gram’s Iodine test was carried out by adding 4 mL of Gram’s solution (VWR) to each plate to cover the colonies and set at room temperature for one minute.

Colonies with no halo around them have disrupted amyE’ and are the colonies that were picked for further cloning/analysis. After inoculating the colonies for downstream applications, a 1 μL sample was taken for culture PCR (see method below) to ensure that the desired circuit length was inserted.

### Strains, Growth, and Fluorescence Characterization

All measurements were conducted using *Bacillus subtilis substr.* 168 PS832 cells containing genome integrated constructs. For each variant, isogenic colonies were used to inoculate overnight cultures in 500 μL of LB media supplemented with 5 μg/mL chloramphenicol (Cm5) in a 96-well deep-well plate. Cultures for characterization were then prepared by a 1:100 dilution using defined M9 media supplemented with 2% glucose and Cm5. Varying concentrations of TNT were prepared from a 1 mg/mL stock solution (Sigma Aldrich) and added to wells at the reported concentrations. Plates were incubated at 30°C with 120 RPM orbital shaking inside a Spark spectrophotometer (TECAN). The OD600 absorbances were recorded every 10 min until the OD600 reached 0.15 to 0.2. Subsequent 96-well microtiter plates were then prepared by a 1:100 dilution of the cultures from the first plates into pre-warmed fresh media (same composition) to continue growing cells. Altogether, cells were maintained in the exponential growth phase for over 12 hours, reaching a steady-state condition. Cells were then sampled at an OD_600_ of 0.15-0.2 from the second plate and fixed in 1X PBS with 2 mg/mL kanamycin. Single-cell mRFP1 fluorescence levels were measured using a BD LSR Fortessa flow cytometer with a green laser (561 nm) and the PE-Texas-Red filter. A total of 100,000 gated events were recorded. Measurements were filtered to remove non-cell events. All fluorescence distributions contained single peaks. The arithmetic means of fluorescence distributions were calculated for each biological replicate. The autofluorescence levels of *Bacillus subtilis substr. 168* cells with no plasmid were subtracted to obtain the final reported mean fluorescence values.

### Stability Assay and mRNA measurements using RT-qPCR

The stability assay was carried out using the same characterization method above, but with a few minor changes. Selected strains were grown with and without TNT (35 µM) exposure and maintained in exponential growth phase (OD_600_ of 0.25-0.3, see Supplementary Data) for approximately 16 hours (day “0”) via serial dilution using LB media supplemented with Cm5. Wild-type *B. subtilis* cells were grown similarly, without antibiotic, as a control. Once cells reached exponential phase, fresh cultures were prepared by a 1:100 dilution from the “day 0” cultures into LB media supplemented with antibiotic, but without TNT. Cells were then maintained in the exponential growth phase for at least 8 hours and then sampled (“day 1”), using 1X PBS with 2 mg/mL kanamycin to fix cells. mRFP1 reporter levels were measured using flow cytometry.

mRNA level measurements were performed on selected strains. Starting from overnight cultures, inoculant cultures were grown in 5 ml Cm-supplemented LB media at 30°C with 300 RPM shaking. Once cells reached exponential growth (OD_600_ of 0.25-0.3), samples were taken to measure initial mRNA levels (timepoint t_0_) and to inoculate cultures with different conditions (adding 35 µM TNT in acetonitrile or adding 0 µM TNT with an equivalent amount of acetonitrile). Cell samples were then extracted after 5 minutes of growth for mRNA level measurements. Cell samples were immediately lysed, followed by total RNA extraction using the Total RNA Purification kit (Norgen Biotek) and DNA removal using the Turbo DNAse kit (Ambion). Following extraction, cDNA was synthesized using the High-Capacity cDNA Reverse Transcription kit (Applied Biosystems) and used as the template for the SYBR Green-based qPCR reaction using the Luna Universal qPCR Master Mix (NEB). The reaction and analysis were conducted using an ABI Step One real-time thermocycler (Applied Biosystems). The housekeeping gene for *B. subtilis*, *gyrA*, was used as an internal control to normalize mRNA level measurements. Fold-changes in mRNA level were calculated using the ΔΔC_T_ method. Primers and measurements are listed in Supplementary Data.

### Characterization of Microbes in Soil Systems

Soil was collected from the organically managed portion of the Russell E. Larson Agricultural Research Farm (40°42’53.0"N 77°55’51.2"W). This soil was a homogenized mixture of soils from multiple plots within the farm and had the following characteristics: 6.9 ± 0.1 pH, P (Phosphorous) ∼30-60 ppm, K (Potassium) ∼100-200 ppm, Mg (Magnesium) ∼100-200 ppm, Ca (Calcium) ∼1000-1500 ppm, organic matter ∼2-4 %, and Total nitrogen levels ∼0.1-0.15. Natural soil was prepared by removing large stones or debris that could impact soil volume, followed by sieving to 2 mm to increase homogeneity. Non-sterile natural soil was used for all experiments unless otherwise noted. When needed for preliminary testing, sterile soil was prepared by autoclaving natural soil for a total of 3 sterilization cycles. Starting from a stock TNT solution (1.2 mg/mL in acetonitrile, Sigma-Aldrich), working TNT solutions were prepared at 0, 100, 1000, and 2500 μM concentrations, using a mixture of acetonitrile and DMSO as solvent. Equal volumes of working solution were applied to aliquots of 10 grams of soil, followed by mixing until saturation. Soil aliquots were then left in a biosafety cabinet to allow for evaporation of the solvent mixture. TNT-treated soil was then combined with 25 grams of untreated soil, mixed thoroughly, and transferred to soil containers.

Nutrient supplementation was used to increase the persistence of the engineered bacteria, based on preliminary soil testing (**Figure S8**). A sugar mixture containing equimolar amounts of fructose, maltose, ribose, glucose, galactose (1:1 ratio by mass) was added at 6 grams sugar / kg soil. A urea solution was added at 100 mg/kg soil. The nutrient mixture was prepared by dissolving in 5 mL deionized water per soil container, followed by filter sterilization (0.2 μm filter) and pipetting to evenly distribute across soil containers. Nutrient supplementation was added once to each soil container.

Engineered *Bacillus subtilis* and control strains were added to soil containers with the following protocol. Three 4-liter sterilized flasks were filled with 2 liters of autoclaved LB media. Each flask was inoculated from an overnight culture, starting from a single colony on a freshly streaked plate. All cultures were incubated at 30°C and 150 RPM shaking for ∼20 hours. Cultures were harvested using a T-1-P Laboratory continuous flow centrifuge (Sharples) or a tabletop centrifuge. Pellets were washed, resuspended with sterile de-ionized water, and their cell densities measured. Resuspended cells were aliquoted to soil containers via pipetting, adding about 10^8^ colony forming units / g soil. For the no cell control, the volume of cell suspension was replaced with the same volume of sterile water. Soil containers were wrapped in parafilm to mitigate dehydration and were periodically re-hydrated to maintain water saturation between 80-85%. The day-0 masses of soil containers were measured using a mass balance. Mass balance measurements were used to track water loss and to determine when rehydration was needed.

Non-destructive soil sampling occurred every 3 to 4 days for a total of 28 days. Samples were extracted in various sections of the plates to ensure homogeneity. For qPCR and NGS, two 250 mg samples were extracted for each timepoint. Genomic DNA was extracted using a NucleoSpin Soil Kit (Macherey-Nagel). SYBR Green PCR Master Mix (Thermo Scientific) was used for quantitative PCR, using gene-specific primers. As a calibration control, qPCR was carried out known concentrations of mRFP1-containing plasmid, using the same gene-specific primers, to create a standard curve that relates ΔC_T_ values to the number of DNA copies per mL (measurements in Supplementary Data). To sequence the genome-integrated synthetic genetic circuit, PCR was carried out on extracted genomic DNA, using primers that amplify the circuit region, followed by nanopore sequencing (Plasmidsaurus).

For flow cytometry measurements and colony counting, three 500 mg samples were extracted. For colony counting, cell-soil solutions were prepared by adding 500 µL LB media to 500 mg soil samples and vortexing. Dilutions were then spread over LB agar plates (with or without 5 µg/mL chloramphenicol as noted), using dilution factors of 10^5^ or 10^6^, followed by incubation at 30°C overnight. Colonies were counted to determine colony forming units per mL cell-soil solution. For flow cytometry measurements, cell-soil solutions were first centrifuged at 4600 RPM and supernatant was decanted, while being careful not to remove cells. 500 µL PBS was then added to each tube, mixed, and stored at 4°C for at least 4 hours to allow soil particles to settle. 20 µL cell samples were then combined with 180 µL PBS, supplemented with 2.2 mg/mL kanamycin, to fix cells. Flow cytometry was then carried out using a green laser (561 nm) and the PE-Texas-Red filter. Distinct cell populations were identified according to their forward scatter and side scatter clusters (see **Figure S11**). Red fluorescence levels were recorded for all cell events in each gated cell population. Engineered *B. subtilis* cells were identified and counted by their clustering in FSC-SSC space as well as their higher red fluorescence levels. mRFP1 fluorescence levels were determined by subtracting the auto-fluorescence of wild-type cells (*B. subtilis* and natural microbes) in soil samples from measured red fluorescence levels. mRFP1 activation ratios in soil are the mRFP1 fluorescence levels of engineered *B. subtilis* cells grown in soil with added TNT divided by the mRFP1 fluorescence levels of engineered *B. subtilis* cells grown in soil without TNT. The fractions of gated cell populations were quantified by dividing the number of gated events by the number of total events.

## Statistical Analysis

For statistical pairwise comparisons, we used two-tailed, two-sample t-tests to determine p-values with 95% confidence levels (α = 0.05). All sample means, standard deviations, replicate numbers, and p-values are provided in Supplementary Data.

## Data Availability

All genetic part and system sequences, model calculations, experimental measurements, and statistical & data analysis are available in Supplementary Data.

## Contributions

T.H.B and H.M.S conceived the study. E.A.E designed and carried out experiments with assistance from G.E.V, D.P.C. and E.G. E.A.E and H.M.S performed data analysis and wrote the paper.

## Acknowledgements

This project was supported by funds from the Defense Advanced Research Projects Agency (HR00111920021) and the National Science Foundation (MCB-2131923).

