## Supplementary Information for "An Autonomous Microbial Sensor Enables Long-term Detection of TNT Explosive in Natural Soil"

#### Supplementary Notes

##### Developing Explanatory Model to Observe Circuit Behavior

To understand the bioreporter's behavior when sense and antisense transcription is varied, we developed an explanatory, mechanistic model using a system of equations to capture mRNA production and monitor the changes in translation rate of the riboswitch as sense and antisense transcription is varied. We started with a simple, classic model for mRNA of a single gene [1], and then accounted for transcriptional interference and polymerase collisions. First, we solved for the RNA polymerase concentrations being produced from the forward promoter with colliding antisense promoters using a transcriptional interference model developed to explore the RNA polymerase collisions between forward promoters ($C_F$) and antisense promoters ($C_R$) [2]. This model used a set of differential equations that solved for polymerase concentrations in both the forward and reverse directions as a function of the length of DNA between promoters, the probability of polymerases dissociating, and the predicted firing rates of the promoters (Equation S1, 2). The differential equations used in the Brophy model capture the effect of collisions in polymerase concentrations of forward and reverse transcripts as a function of DNA length:

$$\frac{dC_F}{dx} = -\varepsilon_F C_F C_R \quad (1)$$

$$\frac{dC_R}{dx} = -\varepsilon_R C_F C_R \quad (2)$$

where the $\varepsilon_F$ and $\varepsilon_R$ parameters represent the frequency of RNA polymerase encounter collisions and dissociate from the DNA, and their values set based on experiment results in Brophy et al. The boundary conditions are the rates that RNA polymerases being elongating, where $C_F(x=0) = \phi_F / v$ and $C_R(x=N) = \phi_R / v$, $\phi_F$ and $\phi_R$ are the promoter firing rates, $v$ is the elongation rate (held constant at 75 nts/s for *B. subtilis* [3]), and N is equal to the length of DNA being transcribed (1709 nts). To calculate the promoter firing rates for the selected promoters in our system, the sequences were individually inserted into the Promoter Calculator to get a predicted transcription rate (au), then a design feature of the calculator was used to convert the scale of relative units to absolute units (RNAP/DNA/s) [4]. To imitate the conditions for circuits tested in Figure 3, which are integrated in *B. subtilis*, we set the copy number to 3, which is ideal scenario based on the amyE' integration location of our circuit [5]. The we calculated the input firing rates for the model with units of RNAP/sec. Using the model and our inputs, we obtained the polymerase concentration per nucleotide (RNAP/nt) for each forward transcript, $C_F$ (x = N), and reverse transcript, $C_R$ (x = N/2), during interfering collisions. Here, the $C_R$ (x = N/2) term represents the polymerase concentration of the reverse promoter at half the length of the DNA segment, which is the assumed location of RNAP collisions.

The resulting polymerase concentrations were then used to predict the amount of mRNA formed from transcriptional regulation between various sense and antisense promoter combinations. To do this, we developed two equations that capture the effect that transcriptional interference and RNA polymerase collisions have on mRNA levels as promoters are varied (Equation S3, 4). During polymerase collisions of competing promoters, double stranded RNA forms when transcripts, expressed at lower levels, pair with their antisense transcripts for downstream targeted degradation [6]. We captured the rate of formation of dsRNA from the collisions of the forward and antisense promoters using the following equation,

$$\frac{dm_{dsRNA}}{dt} = \left[\min\left(C_F(x=N), C_R\left(x=\frac{N}{2}\right)\right) * v\right] - \lambda_{dsRNA}(m_{dsRNA}) \quad (3)$$

where mdsRNA represents the mRNA concentration of the dsRNA formed during antisense transcription collisions, $\lambda_{dsRNA}$ is degradation rate of dsRNA with a 30 second half-life (0.023 $s^{-1}$), v is the elongation rate, and min($C_F$(x=N),$C_R$(x=N/2)) represents the lowest concentration of polymerase from collision interference that will form dsRNA.
The rate of mRNA formed between the sense and antisense promoters, can then be modeled with the following equation,

$$\frac{dm_F}{dt} = [C_F(x=N) * v] - \lambda_{mRNA}(m_{dsRNA} + m_F) \quad (4)$$

where mF represents the mRNA concentration of the DNA being transcribed in the forward direction between promoters, $\lambda_{mRNA}$ is the degradation rate of mRNA with a 5minute half-life (0.0023 $s^{-1}$), and $v$ is the same elongation rate chosen above. Assuming steady-state conditions, equations 3 and 4 are set equal to zero and combined to solve for $m_F$ for each antisense and sense promoter combination tested (Eq. 5).

$$m_F = \frac{\left[\min\left(C_F(x=N), C_R\left(x=\frac{N}{2}\right)\right) * v\right]}{\lambda_{dsRNA}} + \frac{[C_F(x=N) * v]}{\lambda_{mRNA}} \quad (5)$$

Each mRNA concentration calculated was used as an input into the Riboswitch Calculator to determine the translation rate ($r_{TL}$) of the riboswitch at that specific value. By utilizing the same RNA aptamer sequence and its binding affinity, but exchanging mRFP1 for the Int2 integrase, we used the prediction mode of the Riboswitch Calculator to predict the interactions between the RS14 riboswitch, TNT, mRNA, and the ribosome. Inputs into the model include the partial CDS sequence of the Int2 integrase (60 nts), the molar volumetric ratio of the Int2 integrase and water (10,000), the intracellular concentration of TNT added (μM), and the initial mRNA concentration ($m_F$) of the Int2 gene (μM). Similarly to mRFP1, the molar volume of Int2 ($\sim 10^{-21}$ liters) was estimated using the 30S ribosome as reference [7]. Typically, it is assumed that the intracellular concentration of a ligand is 100-fold less than its added extracellular concentration, but this considers only normal growth conditions and not the toxicity effect on cell growth [7]. We found that in LB media, the doubling time for WT B. subtilis at 35 μM of TNT is 3.5 times longer than that with no TNT (Supplementary Data, Riboswitch Results). By assuming the 100-fold intracellular concentration (0.35 μM) occurs during normal growth conditions, we set it's specific growth rate at 0.023 (1/min). We the estimated the intracellular concentration during 35 μM of TNT growth, with a specific growth rate of 0.0067 (1/min). This resulted in an estimated intracellular concentration of 0.01 μM, which is approximately 350-fold less than the extracellular concentration added. From the model, we extracted the translation rates of the riboswitch in the uninduced states ($r_{TL,0\mu M}$) and the induced states ($r_{TL,35\mu M}$) for all mRNA levels. Translation activation ratios are equal to $r_{TL,35\mu M}$ divided by $r_{TL,0\mu M}$. The translation rate in the uninduced state remained unchanged, since it depended on the total Gibbs free energy change of the structure with no ligand bound [7].

#### Supplementary Tables

**Table S1:** RNA-Seq measurements of the *Bacillus subtilis* transcriptome, showing read counts (TPMs) of selected transcripts when growing cells were exposed to different TNT concentrations alongside the genes' promoter and ribosome binding site sequences.

| **ID** | **Product in *B. subtilis* genome** | **Promoter and RBS sequence** | **Read Counts (no TNT)** | **Read Counts TNT 10 *μ*M** | **Read Counts TNT 66 *μ*M** |
|---|---|---|---|---|---|
| QX56_11955 | ATPase AAA | ACTTGCACACACAACTCCGTTCTCTGTCTGCTCGCTGGAAAGCTGTTTCATTTTATGATAAGCCCCTCTGAGCTTAAAAGATCTGACAACTTGCAAGTCTTCGCGTTTTAAATAGATATTGCATTCATATCTCTCAGAAAGTCTGTCATTCCTCTGCAAAGGTGTGTGAATGACAACGTCCTTTACGTTTTGGTGAGCCTTCAAAATGTCTTTCACTTGGATGAGAGAGTTTTCTTTAAGCAACGGTTTCATGTACAGATTCCTTTCTTGTTTTAAATCCCTATTTAATATGCCATTATAACATGAATATTCAAAAAATAAGACAATTATTTTTATTTAATAATTAAAAAAATACAGACTATGTCCACCACATCTTGTGGTAACTTCTGTTCAACCATTATCCTGCAGAGCCAAGAAAAAAGATGATCGTAAAATAAAGGGTGTTTTCAACCAAAAGGCATGATATAATGAAACCAATTAGAACCAAAAGGAGCCAATTG | 428 | 884 | 4267 |
| QX56_18325 | glycine/betaine ABC transporter ATP-binding protein | CATCAAGCATTTCCCGCACTACCTGGCTCATACGTGTTTTACTCATGCCGGTGGCCTCAGATAATTCCGTAAGCGTCATCGGTTTTCGATTCATATAAATAATGCCCAGCACTCGTCCGACGGTAGAGGGCATTCCAAATGCATGCATGTTTTCCGCGATTCTTTCTATAAAATGGTCCTCGGCTTGTTCAATGATCGTTAACGGATCTTTCTCCAACGGTTTCATCCTTTCAGCTAACAATTCCGATGTCAATATACCACATTTTTGATACGGCGAAAATGCTGTCAAAAGAGAATTTGGCAGGAAATTGGAGCAATAAGGAGATCAACACACGTCAATGCGGGTAAATCAGCGTTTACACATACTTTATAAATAATATGAACAAATTGTAAACTTTTTATTTTATAAACTTTATTCTATAATGGGAAGCATTCAATTGTCTGAAAAATTAAATTTAAACTGAACAAATTGAATAAACTTAATTTTGGAGGTGCGATGT | 11 | 37 | 101 |
| QX56_01665 | hypothetical protein | AGCTCCACGATAATGCCGGTATTCTCATCCTGAGGATCTGCCCAACAAAAATGAGCTCTTTTATCATCACCGCCATAATCTTCAATTTCCAGTTTTAAGTGTGTCGGCACCCGGCCGATTTCCAGTGCTTTTTGTTTCAGTCCCTCTTTCTTCACAAAGGCGGCCCCTTTCATCCTTCTTAAGTTCCCTCCAACTAATCAAAAAGTAATAGTCTAAAATACACAGTTTTCTGAGTTTACCAGTTTTACCAGAACATTTTTCACCAAAAATAGGATGATGAACCTATTCAATATAAAAGCTAATAATCTTAAAATTAATTCCGTTGTAACTTTTTATAGGTCCATTCATGACAAAGGAGGGCGGAGTTTCCAGAAAATCCGGTGAAAACCGTGAGAAAAGAACTGATTTTTTTGGTTATATGATGCTAATCCATTCAAAAAGATCCATTTCTCAAAGACAAAAGACTCCCCTTTCCTAAACCATTTGAGGAGGAAAAATGA | 169 | 310 | 1450 |
| QX56_21305 | amino acid ABC transporter ATP-binding protein | TCCGAGGCGTGCCGACACTTGTACAGCTGTTCTTAATCTATTACGGGCTGCCGCAGCTATTTCCAGAGATGAGCAAAATGACAGCTCTCACAGCTGCCATCATCGGGTTAAGCTTAAAAAACGCAGCTTATTTGGCAGAAATCTTCCGGGCCGCCCTCAATTCTGTTGATGACGGGCAGCTGGAGGCGTGCCTGTCTGTCGGTATGACAAAATTTCAGGCATACAGACGGATTATTTTGCCGCAAGCGATCCGAAATGCGATTCCGGCAACGGGCAATACATTTATCGGGCTCCTGAAAGAAACGTCACTGGCCTTTACATTAGGGGTCATGGAGATGTTCGCCCAAGGGAAGATGTACGCTTCAGGAAACCTCAAATATTTTGAGACGTATTTGGCGGTTGCGATCGTCTATTGGGTTCTTACCATTATCTACAGCATTTTGCAGGACTTGTTTGAACGTGCCATGAGCAAGCCATACCGGACTTAGAGGTGAAATCAG | 93 | 202 | 773 |
| QX56_21310 | ABC transporter permease | GGAAACGTATAACTTTACGAAACCATACGCTTATGCGGGAACACAGATTGTCGTCAAAAAAGACAATACAGACATCAAATCAGTAGACGATTTAAAAGGCAAGACAGTCGCAGCCGTTCTCGGTTCAAACCACGCGAAAAACCTTGAAAGCAAAGATCCTGATAAAAAAATCAATATCAAAACGTACGAAACACAAGAGGGTACGCTGAAGGATGTTGCGTACGGCCGTGTAGACGCTTATGTCAACAGCCGAACTGTATTGATCGCGCAAATCAAGAAGACCGGTTTGCCATTAAAGCTTGCAGGAGATCCGATTGTTTACGAACAGGTTGCATTCCCATTTGCCAAGGACGATGCGCACGACAAGCTCCGCAAAAAAGTCAATAAGGCCCTAGATGAATTGCGTAAAGACGGAACACTGAAAAAACTCTCTGAAAAATACTTTAATGAAGATATCACAGTAGAACAGAAGCATTAAGAAACGGCGGTGACTCACCAGA | 42 | 99 | 339 |

### Supplementary Figures

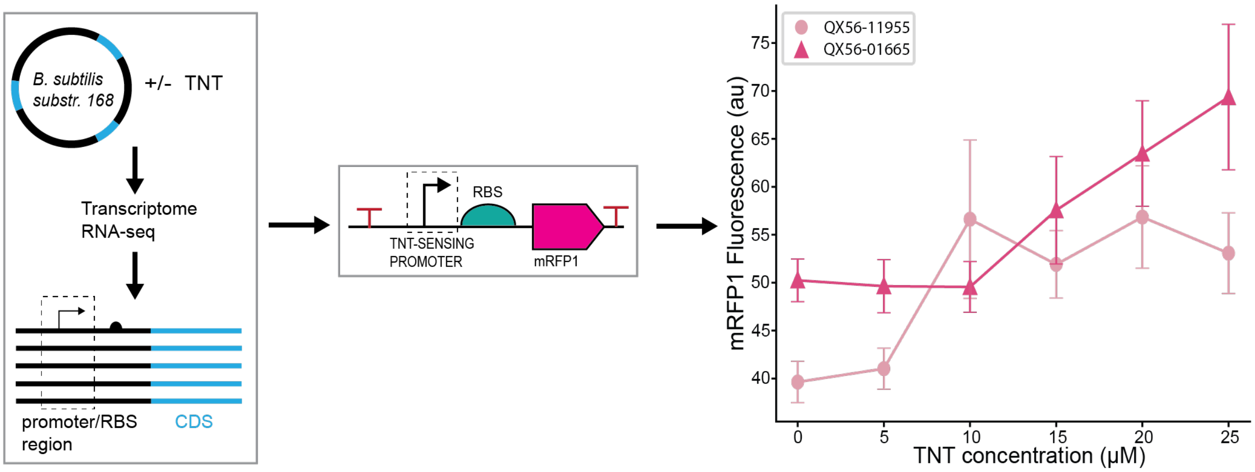


**Figure S1: Design and testing of candidate TNT-sensing promoters in *Bacillus subtilis*.** The *Bacillus subtilis* 168 PS382 strain was cultured in different media conditions (with or without TNT) to reach steady-state mRNA levels. The mRNA was extracted and whole transcriptome RNA-seq was performed to identify genes with elevated mRNA levels (up to 10-fold, see **Supplementary Table 1**) in response to 0.066 mM TNT as compared to the no TNT control (left). The promoter regions from the identified genes were inserted upstream of a designed RBS (25,000 au), mRFP1 reporter protein coding, and transcriptional terminator, followed by integration into the *B. subtilis* genome at the *amyE* locus (middle). mRFP1 fluorescence levels of the TNT-sensing promoters, QX56-11955 and QX56-01665, in response to varied concentrations of TNT (right). Data points and error bars are the mean and standard deviation of N = 3 biological replicates.

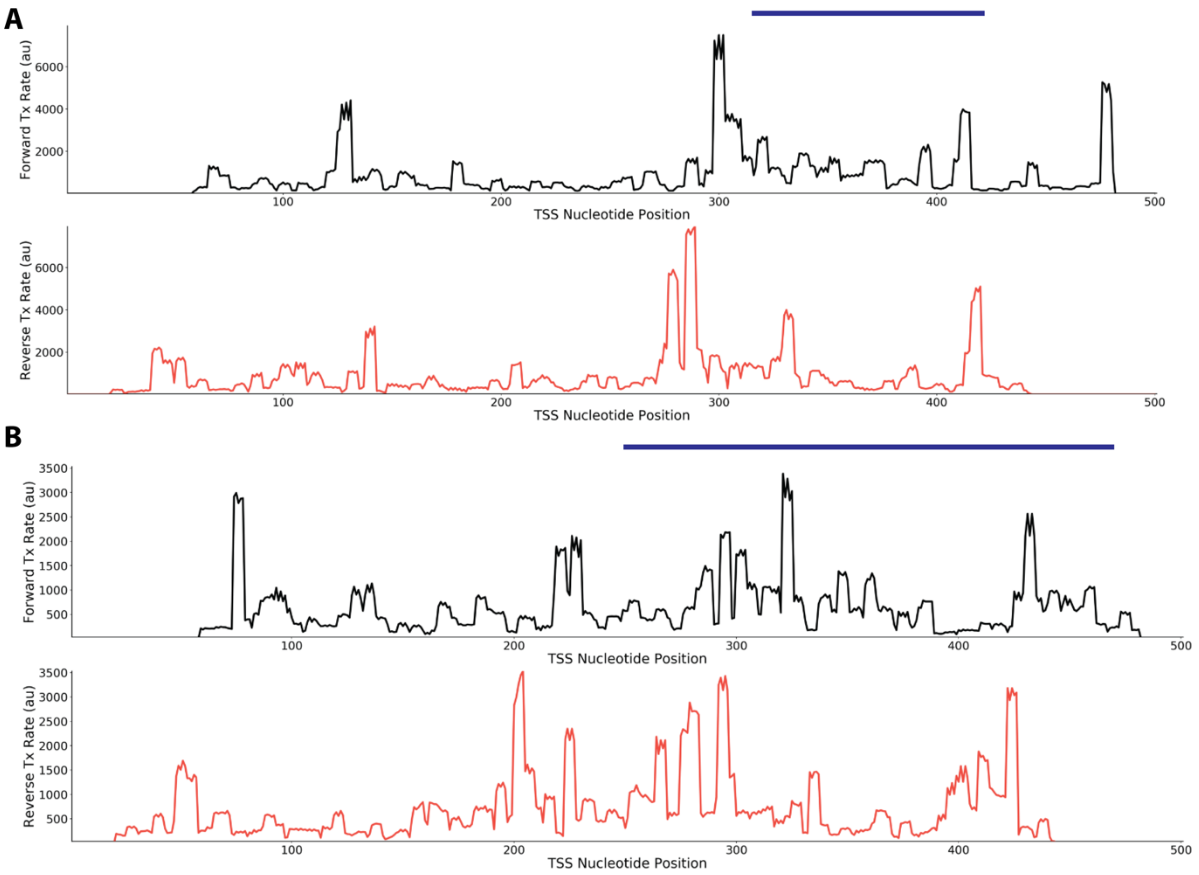


**Figure S2:** Predicted site-specific transcription initiation rates (forward and reverse directions) for two promoter regions corresponding to two genes (**A**: QX56_11955; **B**: QX56_01665) identified as TNT-responsive according to transcriptome RNA-Seq measurements. Blue bars indicate the promoter regions extracted for characterization in a synthetic genetic circuit controlling mRFP1 expression when integrated into the *B. subtilis* genome. Predictions were made using the Promoter Calculator v1.0.

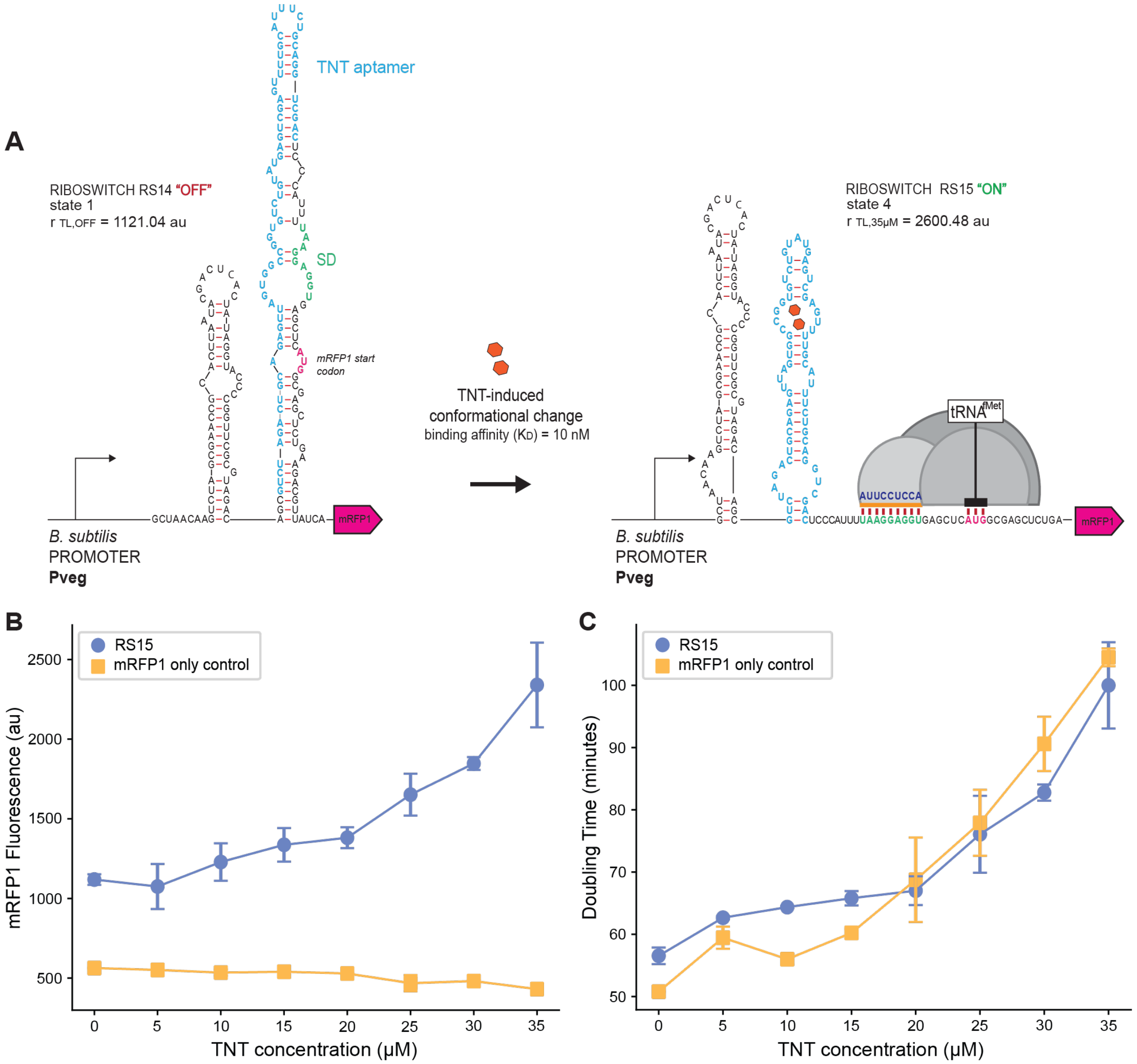


**Figure S3: Design and function of a second TNT riboswitch, RS15, in *Bacillus subtilis*.** **(A)** The Riboswitch Calculator model predicted mRNA structures of the RS15 riboswitch before and after induction with 35 μM of TNT. The "OFF" state (state 1) shows the mRNA structure and translation initiation rate calculated when TNT is unbound ($r_{TL,OFF}$). The "ON" state (state 4) shows the change in mRNA structure and translation rate when both TNT is fully bound to its aptamer and the ribosome is bound to the mRNA ($r_{TL,35\mu M}$). Nucleotides are color coded based on their interactions, (light blue) the TNT aptamer sequence, (green) the Shine-Dalgarno sequence (SD), (dark blue) the last 9 nucleotides of the 16S ribosomal RNA, and (pink) the start codon for mRFP1. The orange bar is the ribosomal footprint for initiation. For riboswitch characterizations, we added a strong *B. subtilis* promoter upstream to measure **(B)** the mRFP1 fluorescence levels and **(C)** the growth rates (doubling time in minutes) of the TNT-RS15 riboswitch strain (blue dots) and mRFP1-only control (yellow squares) in response to varied concentrations of TNT, up to 35 μM. Data points and error bars are the mean and standard deviation of N = 3 biological replicates.

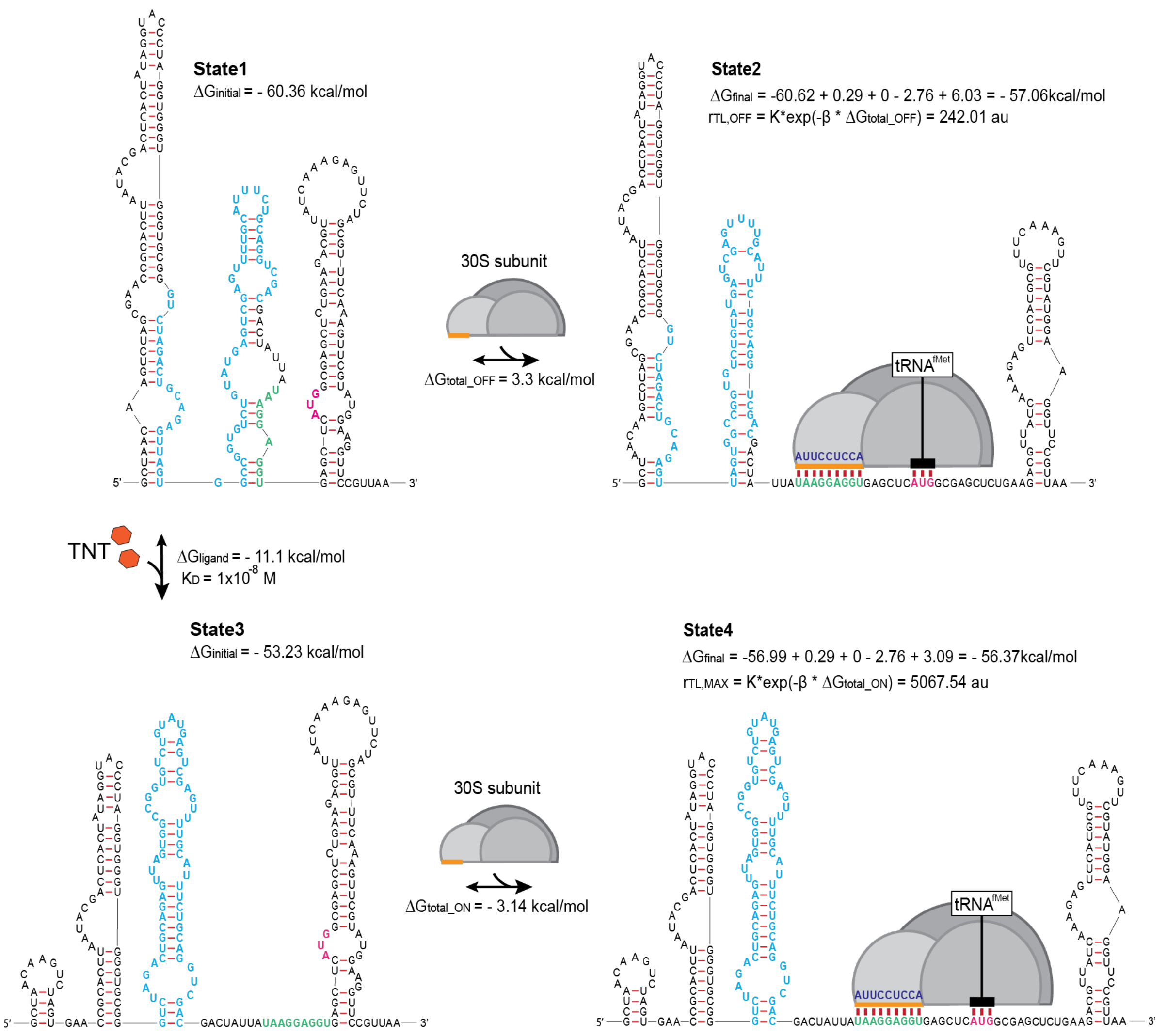


**Figure S4: The Riboswitch Calculator predicted mRNA structures and ribosome-binding free energies for the RS14 riboswitch.**

State 1 shows the initial mRNA structure in the absence of TNT with free folding energy, $\Delta G_{initial}$. State 2 shows the changes in mRNA structure when the ribosome binds to the mRNA in State 1. The free energy model for calculating $\Delta G_{final}$ includes the Shine-Dalgarno (SD) sequence binding to the last 9 nt of the 16S ribosomal RNA ($\Delta G_{mRNA\text{-}rRNA} = -60.62$), binding between the $tRNA^{fMet}$ and the start codon ($\Delta G_{start} = -2.76$), and the energetic penalties for non-optimal spacing ($\Delta G_{spacing} = 0.29$) and an upstream standby site ($\Delta G_{standby} = 6.03$). State 3 shows the change in mRNA structure when TNT binds to its RNA aptamer with a binding free energy of $\Delta G_{ligand}$, which results in a change of free folding energy ($\Delta G_{initial} = -53.23$). State 4 shows the change in mRNA structure and final Gibbs free energy ($\Delta G_{final}$) when the ribosome binds to the mRNA in State 3. The translation rates, $r_{TL,OFF}$ and $r_{TL,MAX}$, are predicted based on their total Gibbs free energy between final and initial states ($\Delta G_{total}$), according to Boltzmann's relationship [7]. Light blue nucleotides represent the TNT aptamer sequence, green represents the Shine-Dalgarno sequence (SD), dark blue represents the last 9 nucleotides of the 16S ribosomal RNA, and pink represents the start codon for mRFP1. The orange bar is the ribosomal footprint for initiation.

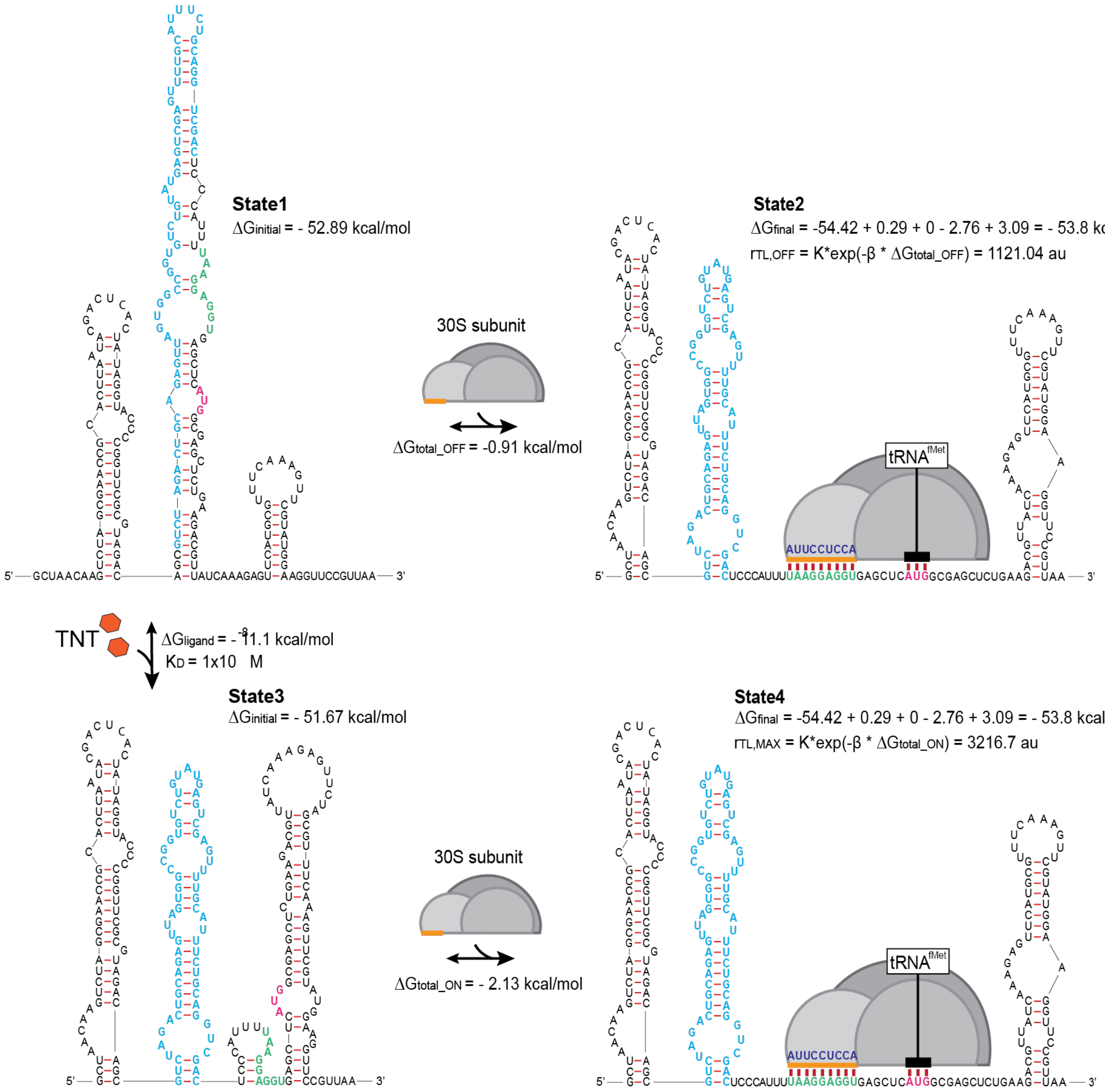


**Figure S5: The Riboswitch Calculator predicted mRNA structures and ribosome-binding free energies for the RS15 riboswitch.**

State 1 shows the initial mRNA structure in the absence of TNT with free folding energy, $\Delta G_{initial}$. State 2 shows the changes in mRNA structure when the ribosome binds to the mRNA in State 1. The free energy model for calculating $\Delta G_{final}$ includes the Shine-Dalgarno (SD) sequence binding to the last 9 nt of the 16S ribosomal RNA ($\Delta G_{mRNA\text{-}rRNA}$ = -54.42), binding between the $tRNA^{fMet}$ and the start codon ($\Delta G_{start}$ = -2.76), and the energetic penalties for non-optimal spacing ($\Delta G_{spacing}$ = 0.29) and an upstream standby site ($\Delta G_{standby}$ = 3.09). State 3 shows the change in mRNA structure when TNT binds to its RNA aptamer with a binding free energy of $\Delta G_{ligand}$, which results in a change of free folding energy ($\Delta G_{initial}$ = -51.67). State 4 shows the change in mRNA structure and final Gibbs free energy ($\Delta G_{final}$) when the ribosome binds to the mRNA in State 3. The translation rates, $r_{TL,OFF}$ and $r_{TL,MAX}$, are predicted based on their total Gibbs free energy between final and initial states ($\Delta G_{total}$), according to Boltzmann's relationship[7]. Light blue nucleotides represent the TNT aptamer sequence, green represents the Shine-Dalgarno sequence (SD), dark blue represents the last 9 nucleotides of the 16S ribosomal RNA, and pink represents the start codon for mRFP1. The orange bar is the ribosomal footprint for initiation.

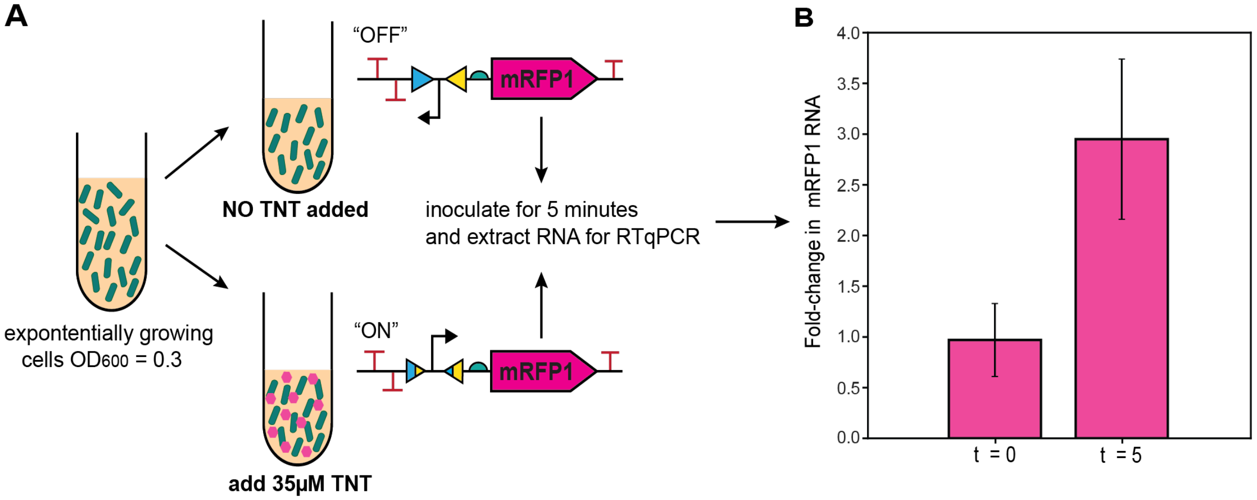


**Figure S6: Measurement of mRFP1 mRNA levels before and after TNT induction. (A)** A schematic of the experimental workflow. The TNT-sensing autonomous microbial sensors growing in liquid culture were exposed to TNT for 5 minutes, followed by sampling, RNA extraction, and RT-qPCR to quantify mRFP1 mRNA levels. The same workflow was carried out on control cells from the same starting culture. **(B)** The fold-change in mRFP1 mRNA levels after 5 minutes of TNT exposure.

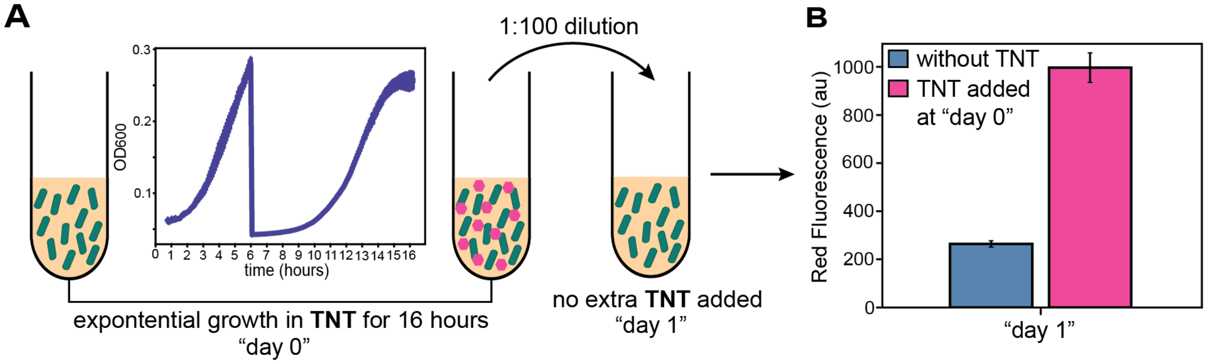


**Figure S7: Characterization of Circuit Functional Stability (Hysteresis) after Removal of TNT Exposure.** (A) A schematic of the experimental workflow. TNT-sensing autonomous microbial sensors were first grown in liquid culture with 35 μM TNT for 16 hours of exponential growth (two serial dilutions), followed by a 1:100 dilution into fresh media without TNT. (B) Mean mRFP1 fluorescence levels were recorded using flow cytometry after an additional 8 hours of exponential growth in the absence of TNT as compared to control cells never exposed to TNT.

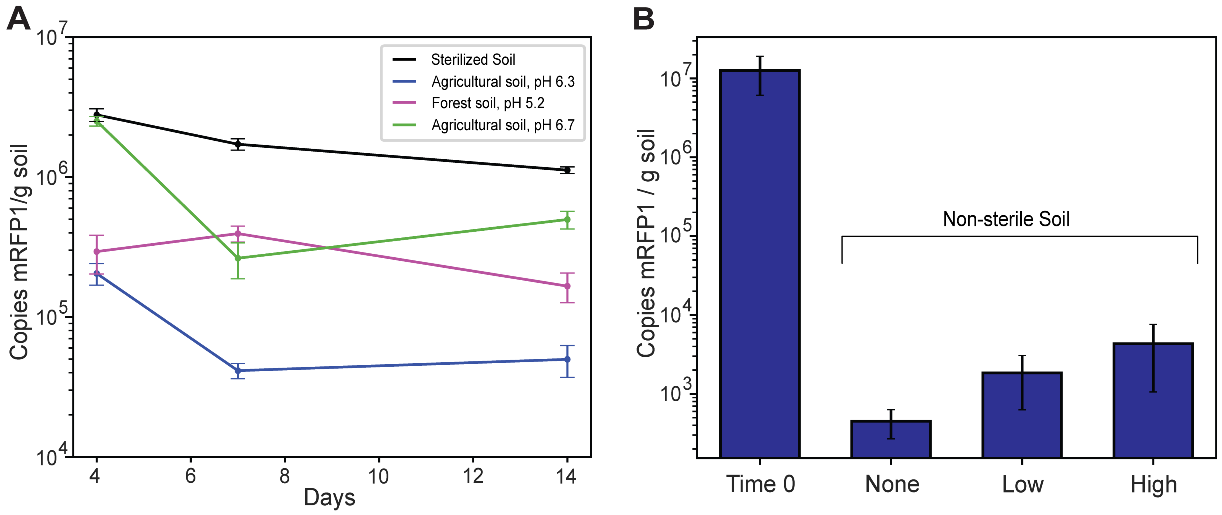


**Figure S8: Effect of pH and nutrient supplementation on genetic performance.**
**(A)** Quantitative PCR measurements of the mRFP1 gene inside soil extractions containing the engineered mRFP1-expressing *B. subtilis* strain across different soil types, (black) sterile soil, (blue) agricultural soil, pH 6.3, (pink) forest soil, pH 5.2, and (green) agricultural soil, pH 6.7. **(B)** Quantitative PCR measurements of the mRFP1-expressing *B. subtilis* strain supplemented with different nutrient additions (none, low, or high) in non-sterile (wild) soil. The same conditions using sterile soil were also tested, however, there was no statistical difference between nutrient additions (Supplementary Data, Preliminary Soil qPCR).

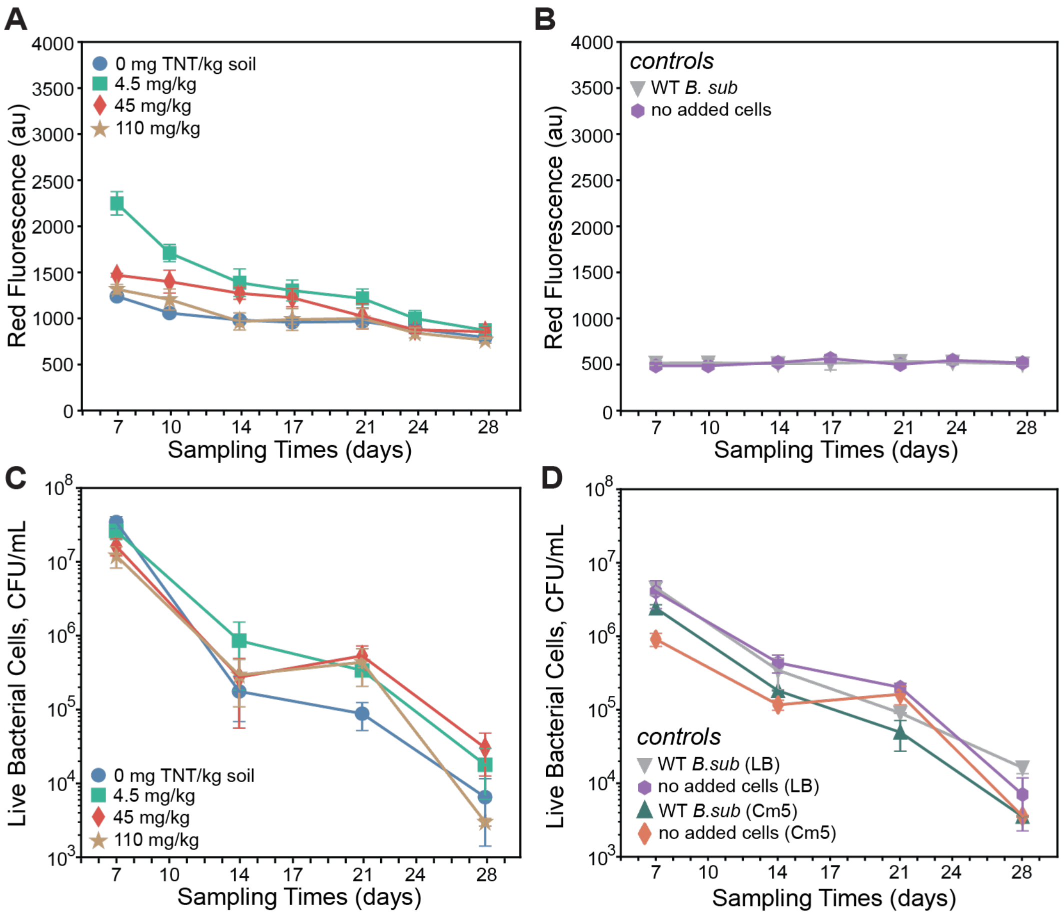


**Figure S9: Long term testing of a TNT "sensor-only" *B. subtilis* strain in wild soil. (A)** Measured red fluorescence levels of the TNT-sensor-only strain in wild soil at 0 mg TNT / kg soil (blue circles), 4.5 mg TNT / kg soil (green squares), 45 mg TNT / kg soil (red diamonds), and 110 mg TNT / kg soil (tan stars) over a 28-day period. **(B)** Measured red fluorescence levels of WT *B. subtilis* cells (grey triangles) and the no added cells control (purple octagons) in wild soil without TNT over a 28-day period. **(C)** Measured cell viability counts on selective agar plates (Cm5) from wild soil samples containing the autonomous microbial sensor with varied TNT amounts. **(D)** Measured cell viability counts on selective (Cm5) or non-selective (LB) agar plates from wild soil samples containing controls. Data points and error bars are the mean and standard deviation of 2 biological replicates (independent soil containers), each with 2 technical replicates (soil samples).

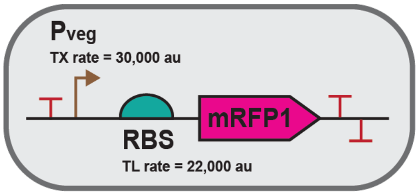


**Figure S10: Schematic of "mRFP1 only" *Bacillus subtilis* strain.**
The "mRFP1 only" strain was constructed using Pveg, a constitutive promoter with a predicted transcription rate of 29,768.4 au (Promoter Calculator v1), and a moderate RBS designed with a predicted translation rate of 22,134.81 au (RBS Calculator v2.1) to express the mRFP1-reporter protein. This circuit was integrated into the B. subtilis genome at the *amyE* locus (1-2 copy number).

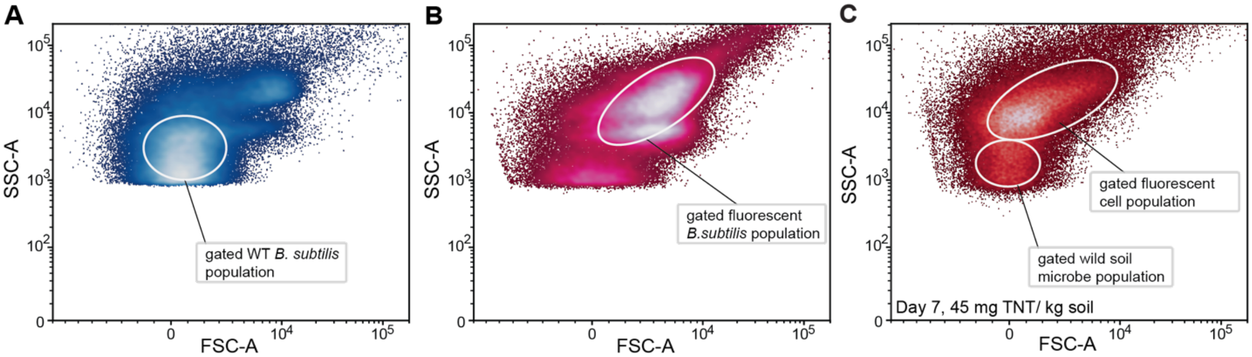


**Figure S11: Gating references of wild type and engineered *Bacillus subtilis* in culture.**
Raw flow cytometry population data (FSC-A vs. SSC-A) and the approximate gating for **(A)** the wild-type *B. subtills substr.* 168 PS832 and **(B)** the engineered *B. subtilis* strain (pD1662:RS14:Int2:#1) after 48 hours of growth in liquid culture, and (C) the engineered *B. subtilis* strain (pD1662:RS14:Int2:#1) after 7 days in "wild" soil inoculated with TNT at a concentration of 45 mg / kg of soil. The gating strategy for **(A)** was FSC-high = $1x10^3$, FSC-low = $-1x10^3$, SSC-high = $7.5x10^3$, SSC-low = $1x10^3$. and for **(B)** was FSC-high = $2.5x10^4$, FSC-low = 0, SSC-high = $1x10^5$, SSC-low = $8x10^3$ and (**C**) shows both gating strategies applied to all soil samples in **Figure 4**.
